# Multi-omic dataset of patient-derived tumor organoids of neuroendocrine neoplasms

**DOI:** 10.1101/2023.08.31.555732

**Authors:** Nicolas Alcala, Catherine Voegele, Lise Mangiante, Alexandra Sexton-Oates, Hans Clevers, Lynnette Fernandez-Cuesta, Talya L. Dayton, Matthieu Foll

## Abstract

**Background:** Organoids are three-dimensional experimental models that summarize the anatomical and functional structure of an organ. Although a promising experimental model for precision medicine, patient-derived tumor organoids (PDTOs) have currently been developed only for a fraction of tumor types.

**Results:** We have generated the first multi-omic dataset (whole-genome sequencing, WGS, and RNA-sequencing, RNA-seq) of PDTOs from the rare and understudied pulmonary neuroendocrine tumors (*n* = 12; 6 grade 1, 6 grade 2), and provide data from other rare neuroendocrine neoplasms: small intestine (ileal) neuroendocrine tumors (*n* = 6; 2 grade 1 and 4 grade 2) and large-cell neuroendocrine carcinoma (*n* = 5; 1 pancreatic and 4 pulmonary). This dataset includes a matched sample from the parental sample (primary tumor or metastasis) for a majority of samples (21/23) and longitudinal sampling of the PDTOs (1 to 2 time-points), for a total of *n* = 47 RNA-seq and *n* = 33 WGS. We here provide quality control for each technique, and provide the raw and processed data as well as all scripts for genomic analyses to ensure an optimal re-use of the data. In addition, we report somatic small variant calls and describe how they were generated, in particular how we used WGS somatic calls to train a random-forest classifier to detect variants in tumor-only RNA-seq.

**Conclusions:** This dataset will be critical to future studies relying on this PDTO biobank, such as drug screens for novel therapies and experiments investigating the mechanisms of carcinogenesis in these understudied diseases.

## Data Description

### Context

Organoids are three-dimensional experimental models that summarize the anatomical and functional structure of an organ [1, 2]. Organoids are revolutionizing fundamental and medical research by allowing us to recapitulate human physiology better than animal models, and also allowing to recapitulate developmental biology contrary to traditional cell cultures [2]. Patient-derived tumor organoids (PDTOs) have been successfully derived for tumors, providing the experimental tools to model disease progression and the preclinical models for personalized treatment testing [3, 4, 5]. Although a promising experimental model, PDTOs have currently been developed only for a fraction of tumor types, focusing on the most frequent cancers and those easiest to culture, leaving rare cancers without appropriate experimental models.

We have recently described one of the very first patient-derived organoid biobanks for the rare and understudied neuroendocrine neoplasms [6]. Neuroendocrine neoplasms are rare tumors that can arise in multiple body sites, predominantly in the lung and gastrointestinal tract [7, 8, 9]. Neuroendocrine neoplasms are further classified into neuroendocrine tumors (NETs) and neuroendocrine carcinomas (NECs). NETs are themselves subdivided into grades (ranging from 1 to 2 or 3 depending on the organs), while NECs are subdivided into small cell and large cell (LCNEC). While small cell carcinomas are more common (e.g., 15% of lung tumors), benefited from more studies and have dedicated treatment options [10], the best treatment option for LCNEC is still unclear [11], and although most NETs progress slowly and have a good prognosis, a subgroup of tumors metastasize and relapse [12].

We report here the multi-omic dataset (whole-genome sequencing, WGS, and RNA-sequencing, RNA-seq) of the neuroendocrine neoplasm PDTO biobank described in [6] (see Table 1). The dataset contains PDTOs of the lung (*n* = 12; 6 grade 1, 6 grade 2) and small intestine ileum (*n* = 6; 2 grade 1 and 4 grade 2), and LCNEC of the lung (*n* = 4) and pancreas (*n* = 1). This dataset includes longitudinal sampling of the organoids (2 to 3 time-points), and sequencing of the matched parental tumor for most samples (21/23, either primary tumors or metastases). Along with raw and processed data, we provide quality controls for each technique and scripts to run a complete molecular analysis. This unique dataset will provide a reference for future research on the understudied neuroendocrine neoplasms.

**Table 1.**
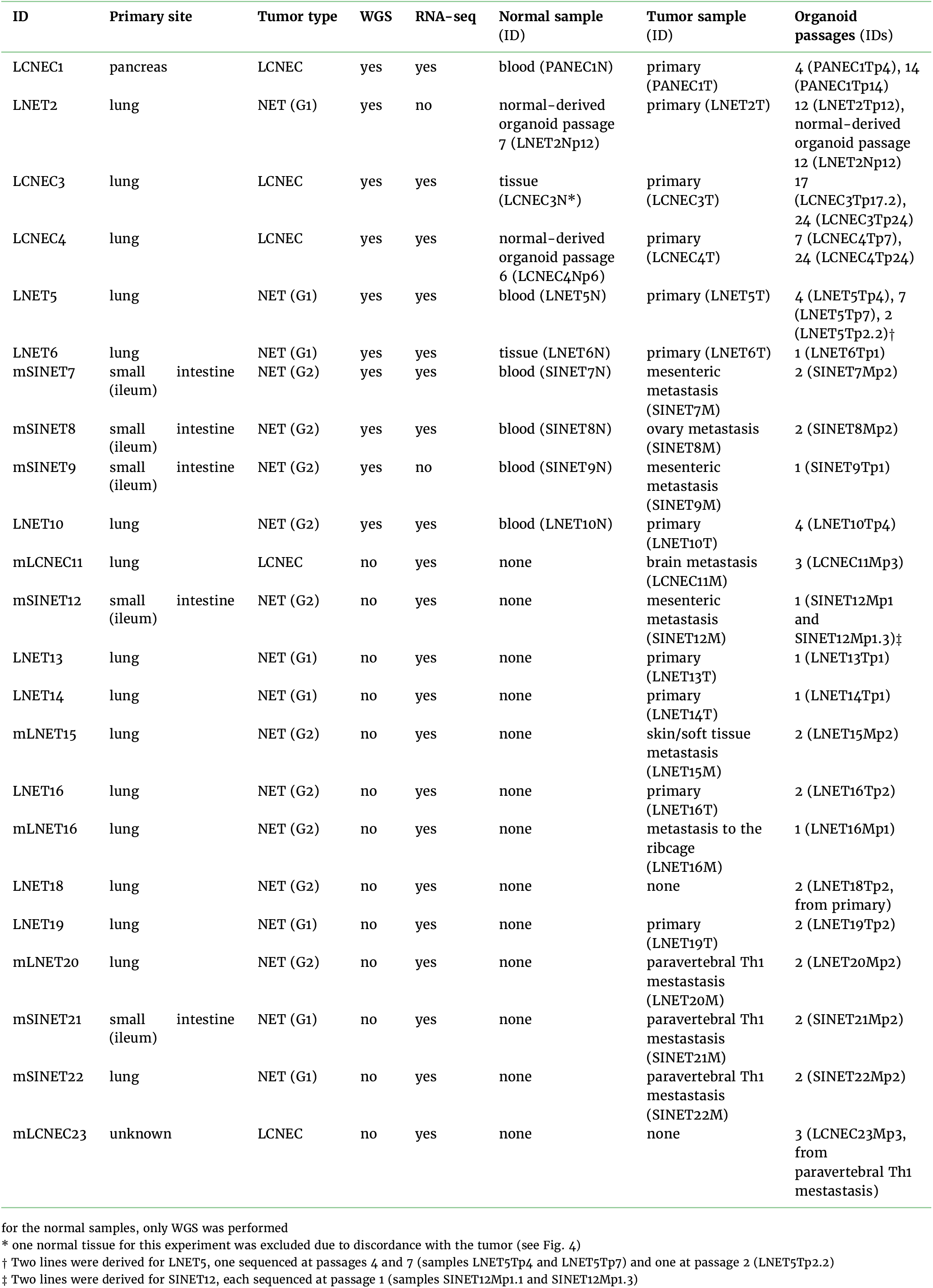
Sample summary.

#### Key Points

- Tumor-derived organoids are revolutionary experimental resources to test biological hypotheses and treatment options
- We have generated the first multi-omic dataset for neuroendocrine tumor organoids of the lung, and for the rare neuroen-docrine tumors of the pancreas, and small intestine (ileum)

## Methods

### Sample collection

PDTO lines of the biobank described in [6] were established from surgical resections or biopsies, put in culture and expanded. PDTOs periodically underwent passaging, a process by which organoids are subcultured to allow future growth [13]; passage time varied from a week to several months depending on the growth rate ([6] Fig. 2). H&E stainings were performed and samples underwent an independent pathological review, and immunohistochemistry of common neuroendocrine markers (Chromogranin A, synaptophysin) were performed to confirm the tumoral neuroendrocrine nature of the parental tumors and PDTOs. See [6] for a detailed description of the protocol.

### Extraction

For each tumor or PDTO, DNA and RNA were extracted from the same sample using the QIAGEN All Prep DNA/RNA Mini kit.

### Sequencing

*Whole-GenomeSequencing (WGS)*. Whole-genome sequencing was performed by the Utrecht Sequencing Facility. After DNA quality control, genomic DNA (0.5–1 µg) was used to prepare the whole-genome sequencing library, using the Illumina TruSeq DNA Nano Kit. Libraries were then sequenced on a Novaseq 6000 platform, as paired-end 150 bp reads, with a target average coverage of 30X for normal samples and 60X to 90X for tumor tissue and PDTOs.

*RNA-Sequencing (RNA-seq)*. RNA sequencing was performed by the Utrecht Sequencing Facility. After RNA quality control, libraries were prepared using the Illumina TruSeq Stranded mRNA polyA Kit. Libraries were sequenced either on a Nextseq 2000 or an Illumina Novaseq 6000, as paired-end 150bp reads.

### Data processing

All data processing was performed using the workflows developed by the rare cancers genomics team of the International Agency for Research on Cancer / World Health Organization (https://github.com/IARCbioinfo/), as detailed in [14] and [15]. The workflows are written in the popular domain-specific language nextflow [16]. All software dependencies are contained in conda environments and containerized with Docker and Singularity (containers available at https://hub.docker.com/ and https://singularity-hub.org/).

*WGS*. Raw reads were mapped to reference genome GRCh38 using workflow *alignment-nf* (https://github.com/IARCbioinfo/alignment-nf, v1.2). This workflow first maps reads (software bwa-mem2 v2.0 [17, 18]), then marks duplicates (software samblaster, v0.1.26 [19]), and finally sorts reads (software sambamba, v0.7.1 [20]).

*RNA-seq*. Raw reads were mapped to reference genome GRCh38 with annotation gencode v33 using the workflow *RNAseq-nf* (https://github.com/IARCbioinfo/RNAseq-nf, v2.4). This work-flow removes adapter sequences (wrapper Trim Galore v0.6.5 [21] for software cutadapt [22]), maps reads (software STAR v2.7.3a [23]), marks duplicated reads (software samblaster, v0.1.25), and finally sorts reads (software sambamba, v0.7.1). Alignments were then post-processed using two workflows to improve their quality. Workflow *abra-nf* (https://github.com/IARCbioinfo/abra-nf, v3.0) performs local realignment using software ABRA2 (v2.22 [24]), and *BQSR-nf* (https://github.com/IARCbioinfo/BQSR-nf, v1.1) performs base quality score re-calibration using gatk (v4.0.5.1 [25]).

*Variant calling from WGS*. Single nucleotide variants were called on all WGS samples using software Mutect2 from GATK4 (v4.2.0.0 [26, 27]) with workflow *mutect-nf* (https://github.com/IARCbioinfo/mutect-nf, v2.2b), as described in [6]. Resulting variant calling format (VCF) files were normalized using bcftools v1.10.2 [28] (workflow https://github.com/IARCbioinfo/vcf_normalization-nf, v1.1) and annotated using ANNOVAR v2020Jun08 (workflow https://github.com/IARCbioinfo/table_annovar-nf v1.1.1). Indels and multinucleotide variants were additionally filtered using the intersection of Mutect2 and strelka2 [29] calls (workflow https://github.com/IARCbioinfo/strelka2-nf v1.2a), in order to reduce false positives that are more frequent in indel calls due to the diiculty of detecting such variants with short reads sequencing.

*Variant calling from RNA-seq*. Variants were called on all RNA-seq samples using software Mutect2 from GATK4 (v4.2.0.0 [26, 27]) with workflow *mutect-nf* (https://github.com/IARCbioinfo/mutect-nf, branch RNAseq), in RNA-seq and tumor-only modes. The RNA-seq mode incorporates a pre-processing step to fix CIGAR strings (removing NDN elements and ensuring that mapping quality 255 is not used as some mappers like STAR can do), and GATK4’s SplitNCigarReads method that splits reads with Ns in their CIGAR string, in order to improve variant calling quality. Resulting variant calling format (VCF) files were normalized using bcftools v1.10.2 [28] (workflow https://github.com/IARCbioinfo/vcf_normalization-nf, v1.1) and annotated using ANNOVAR v2020Jun08 (workflow https://github.com/IARCbioinfo/table_annovar-nf v1.1.1). For samples which also had WGS data, RNA-seq-detected variants were classified as somatic or germline based on the WGS variant calls described above.

for the normal samples, only WGS was performed

## Quality control

For each ‘omic technique, quality controls (QC) of the samples were performed at each step.

### Raw reads

Software FastQC (v0.11.9 [30]) was used to check raw reads quality, and software MultiQC (v1.9 [31]) was used to aggregate the QC results across samples and generate interactive plots; all plots from Figs. 1 and 2 were generated by multiQC from the FastQC outputs. Original MultiQC reports are available in Supplementary Information (Files S1-S4) to allow a free exploration of the QC statistics.

**Figure 1.**
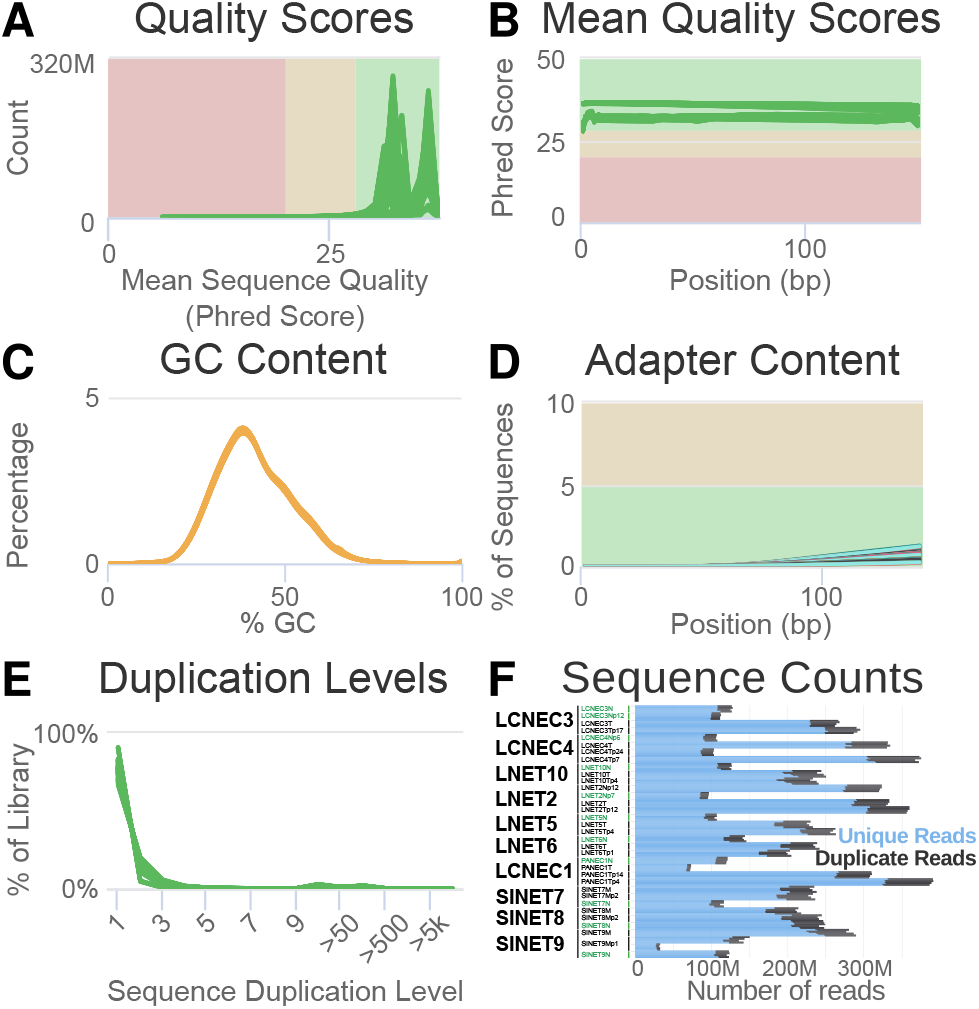
Quality control of the raw Whole-Genome Sequencing (WGS) data. Distribution of the mean sequence quality of the reads in Phred score. (B) Mean sequence quality score as a function of the position in the read in base pairs (bp). (C) Distribution of the GC content in percent. (D) Percentage of reads containing a sequence corresponding to the Illumina adapter sequence as a function of the position in the read in bp. (E) Percentage of the library with a given level of duplication. (F) Number of unique and duplicated reads per file. In panels (A)–(E), each line corresponds to a fastq file, with each of the 34 samples from Table 1 subdivided into four sequencing lanes (except SINET9Mp1, subdivided into 8 lanes), and additionally subdivided into two read pair files, for a total of 4*×*2*×*33+8*×*1 = 280 files; in panel (F), each horizontal bar corresponds to a file. In (A)-(E), green lines correspond to files that passed the most stringent QC filters of software FastQC; orange lines correspond to files that passed a less stringent filter.

**Figure 2.**
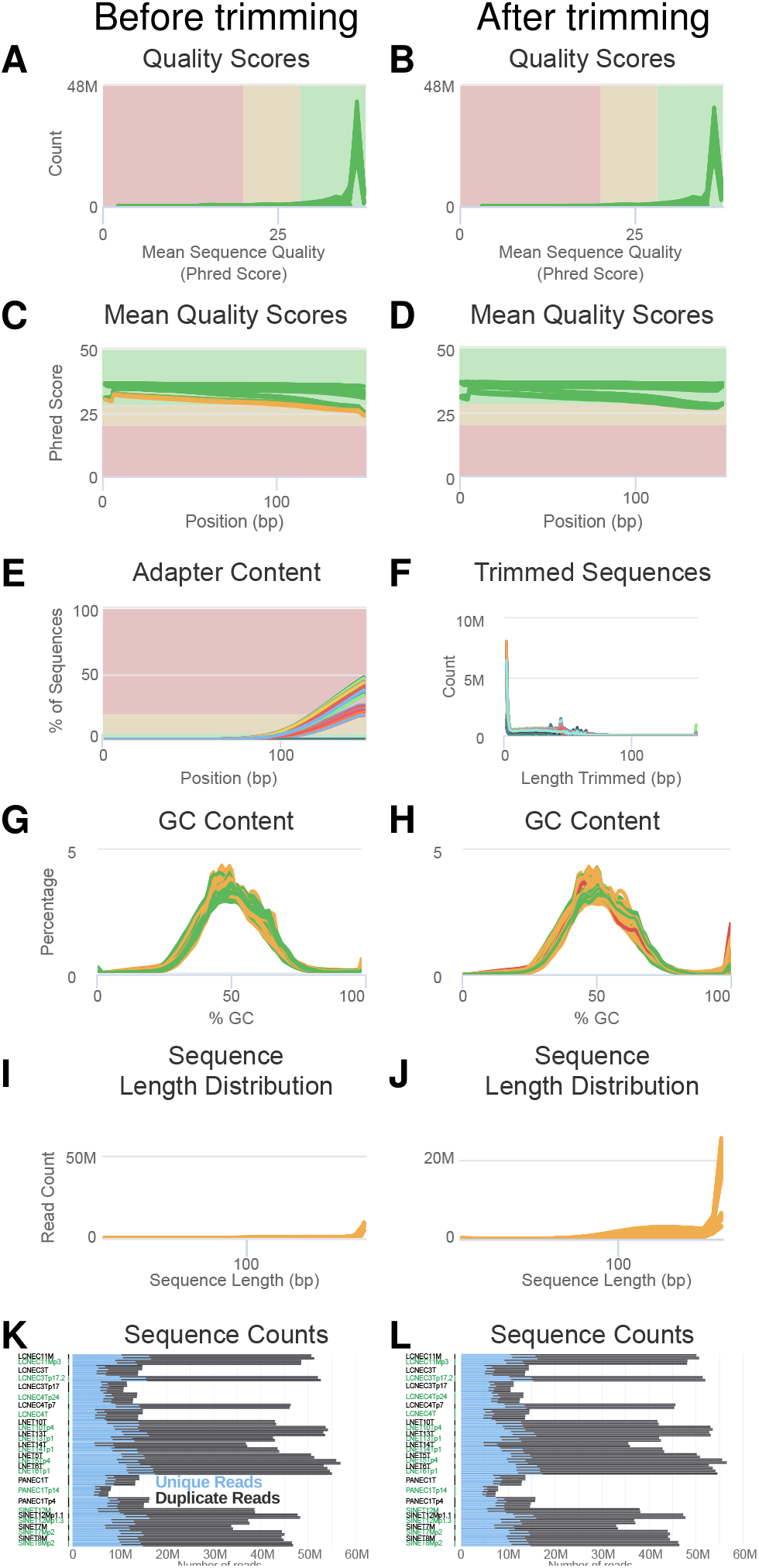
Quality control of the raw RNA-seq data. Panels (A), (C), (E), (G), (I), (K) correspond to controls before read trimming for quality and adapter content by wrapper Trim Galore for software cutadapt; panels (B), (D), (F), (H), (J), (L) correspond to controls after read trimming. Figure legends for panels (A)–(E) and (G)–(L) follow that of Fig. 1. (F) Distribution of the length of the reads trimmed by software cutadapt, for each file (colored lines). In panels (A)–(J), each line corresponds to a fastq file, with each of the 10 non-normal samples from Table 1 divided into two or four sequencing lanes, and further subdivided into two read pair files, for a total of 2 *×* 2 *×* 21 + 4 *×* 2 *×* 7 = 140 files; in panels (K)–(L), each horizontal bar corresponds to a file.

*WGS*. Raw reads passed quality control filters in all samples. All samples displayed good sequence quality scores (mode above 30 Phred), both on average and across all positions in the read (Fig. 1A and B), with samples sequenced later (lower part of Table 1, from LNET5 to LNET10) displaying better scores (highest mode in Fig. 1A). GC content were slightly skewed toward lower values but proved consistent across samples (Fig. 1C), and adapter content and duplication levels were adequate (Fig. 1D and E). The number of reads were consistent between read pairs and consistent with target read depths (Fig. 1F): samples with a target depth of 30X–normal, normal-derived organoids, the primary tumor from experiment LCNEC1, and tumor organoid passage 14 from experiment LCNEC3 (LCNEC3Tp14)–having a lower number of reads (∼ 4 *×* 100M reads= 400M reads) than the others samples (∼ 4 *×* 250*M* = 1000M reads), which had a target depth of 90X. Note that the metastasis organoid of experiment SINET9 (SINET9Mp1) has been sequenced in eight lanes, with 4 lanes with a low number of reads (∼ 30*M*) and 4 additional ones with a larger number (∼ 140*M*) so the total is comparable with that of the other samples.

*RNA-seq*. Raw reads passed quality filters after reads trimming for adapter content and quality. All samples displayed good sequence quality scores on average both before and after read trimming (mode above 30 Phred; Fig. 2A and B), with samples sequenced later (lower part of Table 1, from LNET5 to LNET14) displaying better scores (highest mode in Fig. 1A). Six samples displayed lower scores at the end of the reads before trimming (Fig. 2C) but better scores after trimming (Fig. 2D). Indeed, most samples displayed high adapter content before trimming (Fig. 2E), and the trimming step successfully removed them (less than 0.1% in all samples; Supplementary Information File S1). The trimming step mostly removed less than 5 bp from the read, but occasionally could remove up to around 50 bp (Fig. 2F). GC content were consistent across samples (Fig. 2G-H), although the read trimming step resulted in an excess of reads with high GC content, presumably due to some reads being strongly shortened by the trimming step. Hopefully, in general the trimming step did not increase much the proportion of short reads (Fig. 2I-J). The number of reads were consistent between read pairs and across sequencing runs both before and after trimming (Fig. 2K-L), and total read numbers for each sample were consistent with the target number of 50M (25M pairs): the smallest number, 60.8M corresponded to sample PANEC1Tp14.

### Alignments

*WGS*. The software qualimap (v2.2.2b [32]) was called by our workflow *alignment-nf* to generate QC statistics for the WGS alignments in parallel to the data processing (Table 2). All normal and normal tissue-derived organoids displayed a mean coverage *≥*30X, and all tumor and tumor-derived organoids except passage 24 from the organoid of experiment LCNEC4 (sample LCNEC4Tp24) and passage 1 of the organoid of experiment SINET9 (sample SINET9Mp1) had a coverage *≥* 60X; all samples displayed at least 65% of the genome with a coverage larger than or equal to 30X except LCNEC4Tp24 and (57.4%). Percentages of aligned reads exceeded 99.8% for all samples. Interestingly, some tumor and tumor-derived organoid samples displayed bimodal coverage distributions compatible with variations in copy number state (Supplementary Information File S3).

**Table 2.** Quality control of the WGS alignments.

| Sample Name | % GC | $\geq 30X$ | $\geq 50X$ | Coverage | % Aligned |
| --- | --- | --- | --- | --- | --- |
| PANEC1N | 42% | 84.6% | 16.2% | 41.0X | 99.9% |
| PANEC1T | 42% | 93.8% | 91.7% | 104.0X | 99.8% |
| PANEC1Tp4 | 42% | 93.9% | 93.3% | 127.0X | 99.9% |
| PANEC1Tp14 | 42% | 93.6% | 90.9% | 89.0X | 99.8% |
| LNET2Np7 | 41% | 67.2% | 1.9% | 33.0X | 99.9% |
| LNET2Np12 | 41% | 93.4% | 92.9% | 104.0X | 99.9% |
| LNET2T | 41% | 93.3% | 92.9% | 109.0X | 99.9% |
| LNET2Tp12 | 42% | 93.4% | 93.1% | 115.0X | 99.8% |
| LCNEC3N | 41% | 82.5% | 11.1% | 39.0X | 99.9% |
| LCNEC3Np12 | 42% | 82.5% | 11.7% | 38.0X | 99.9% |
| LCNEC3T | 41% | 93.9% | 90.0% | 89.0X | 99.9% |
| LCNEC3Tp17 | 42% | 93.5% | 90.9% | 90.0X | 99.8% |
| LCNEC4Np6 | 42% | 69.7% | 2.7% | 34.0X | 99.9% |
| LCNEC4T | 41% | 93.0% | 88.1% | 102.0X | 99.8% |
| LCNEC4Tp7 | 42% | 91.7% | 87.9% | 102.0X | 99.9% |
| LCNEC4Tp24 | 42% | 51.9% | 11.3% | 30.0X | 99.9% |
| LNET5N | 41% | 68.9% | 2.5% | 33.0X | 99.9% |
| LNET5T | 42% | 91.9% | 77.6% | 68.0X | 99.9% |
| LNET5Tp4 | 42% | 93.3% | 87.6% | 75.0X | 99.9% |
| LNET6N | 42% | 86.6% | 22.8% | 43.0X | 99.9% |
| LNET6T | 42% | 93.0% | 83.1% | 72.0X | 99.9% |
| LNET6Tp1 | 42% | 90.2% | 76.8% | 61.0X | 99.9% |
| SINET7N | 42% | 77.2% | 4.9% | 36.0X | 99.9% |
| SINET7M | 41% | 92.8% | 83.5% | 73.0X | 99.9% |
| SINET7Mp2 | 42% | 92.8% | 85.7% | 69.0X | 99.9% |
| SINET8N | 42% | 93.1% | 91.7% | 75.0X | 99.9% |
| SINET8M | 41% | 92.6% | 81.2% | 64.0X | 99.9% |
| SINET8Mp2 | 42% | 93.0% | 85.5% | 70.0X | 99.9% |
| SINET9N | 42% | 84.4% | 7.2% | 38.0X | 99.9% |
| SINET9M | 41% | 93.0% | 90.0% | 81.0X | 99.9% |
| SINET9Mp1 | 42% | 90.2% | 49.1% | 49.0X | 99.9% |
| LNET10N | 42% | 86.4% | 9.5% | 39.0X | 99.9% |
| LNET10T | 42% | 93.0% | 90.1% | 71.0X | 99.9% |
| LNET10Tp4 | 42% | 93.0% | 87.1% | 66.0X | 99.9% |

*RNA-seq*. Software RSeQC (v3.0.1 [33]) was called to check alignment quality in parallel to the data processing by work-flow *RNAseq-nf*. For all samples, the number of known junctions (i.e., junctions annotated in the gencode v33 annotation file) was stable when resampling subsets of 75% to a 100% of the reads (all lines plateau in Fig. 3A), indicating a good saturation and suggesting that the sequencing depth was suicient to detect known junctions. In contrast, the number of novel junctions (i.e., junctions not in the annotation file) was increasing slowly as a function of the percentage of reads resampled, but did not completely saturate (no complete plateau in Fig. 3B). This indicates that we probably detected the most abundant novel junctions but that some low abundance novel junctions were probably not detected.

**Figure 3.**
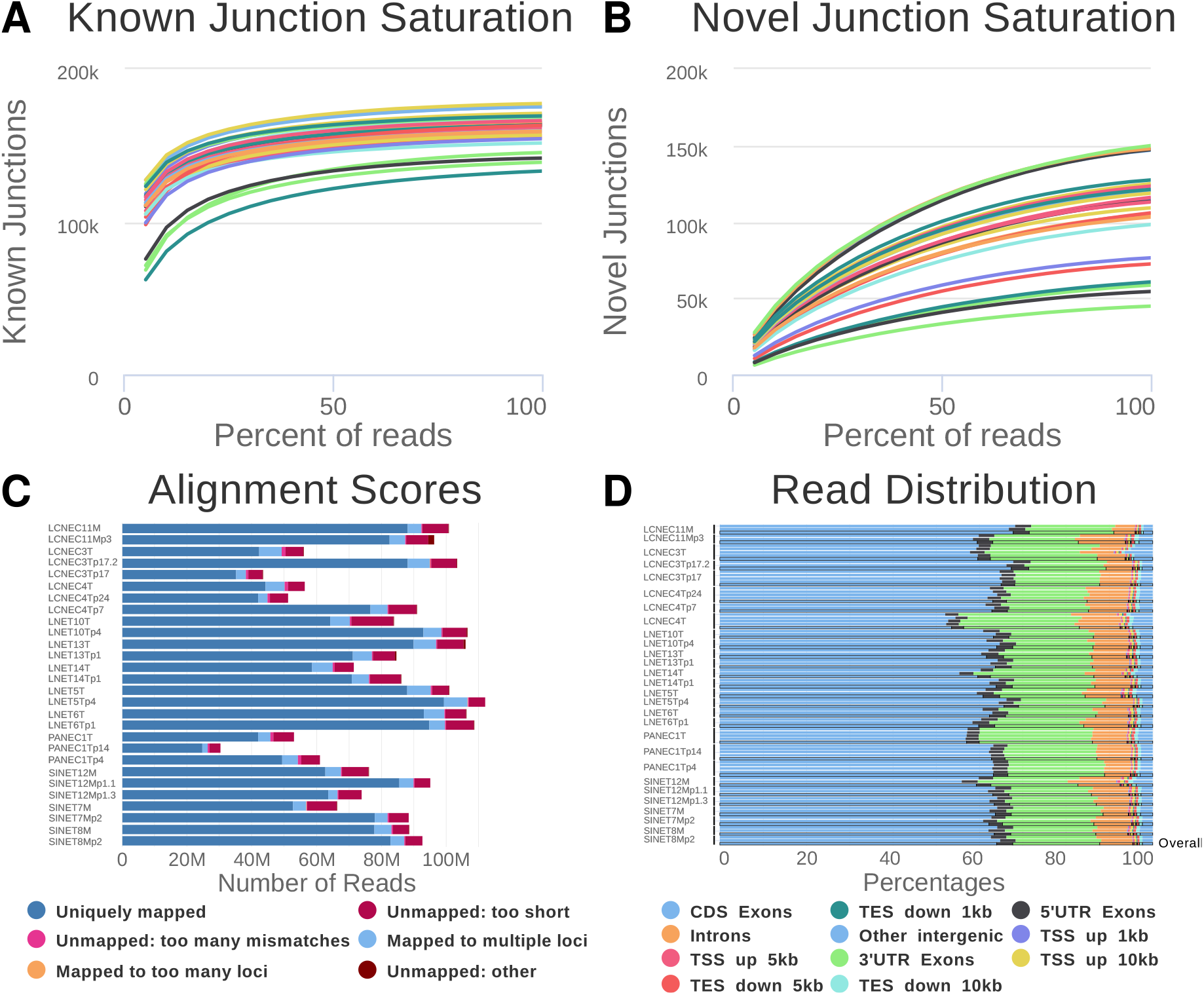
Quality control of the RNA-seq alignments. (A) Number of known junctions identified by software STAR in a subsample as a function of the percentage of reads in the subsample. (B) Number of novel junctions identified by STAR in a subsample as a function of the percentage of reads in the subsample. (C) Number of sequence tags with each alignment score. (D) Distribution of reads among annotated regions.

Alignment scores were good, with more than 25M mapped read pairs (50M reads) for all samples, and from 4M to 7M unmapped reads, mainly due to reads being too short or having too many mismatches (Fig. 3C). The distribution of the alignments within annotated regions matched our expectations, with most reads (*≥*80%) either aligning to exons (*≥*50%), 3’ UTR (∼3%), and 5’ UTR (∼25%) (Fig. 3D).

## Data Validation

### Sample matching

We used software NGSCheckMate (cloned from the github repository https://github.com/parklab/NGSCheckMate revision 10799087bdfe4b990add5b5e536f87c47bbdb688) to check that samples from the same experiment indeed came from the same individual, in both WGS and RNA-seq simultaneously, using our workflow *NGSCheckMate-nf* (https://github.com/IARCbioinfo/NGSCheckMate-nf, v1.1). The sample matching algorithm correctly identified all experiments except one (Fig. 4). The WGS normal-derived organoid sample from experiment LCNEC3 (LCNEC3Np12_WGS in Fig. 4) was found not to match other LCNEC3 samples, suggesting a possible sample swap and was thus excluded from further analyses. Also, the RNA-seq tumor sample for the late-passage organoid of experiment LCNEC3 (sample LCNEC3Tp17_RNA in Fig. 4) was found to better match experiment LNET2, and was thus excluded from the subsequent analyses. Finally, two samples were found to partially match LNET15 and LNET16, suggesting contamination and were also excluded (UNKN00 and UNKN01).

**Figure 4.**
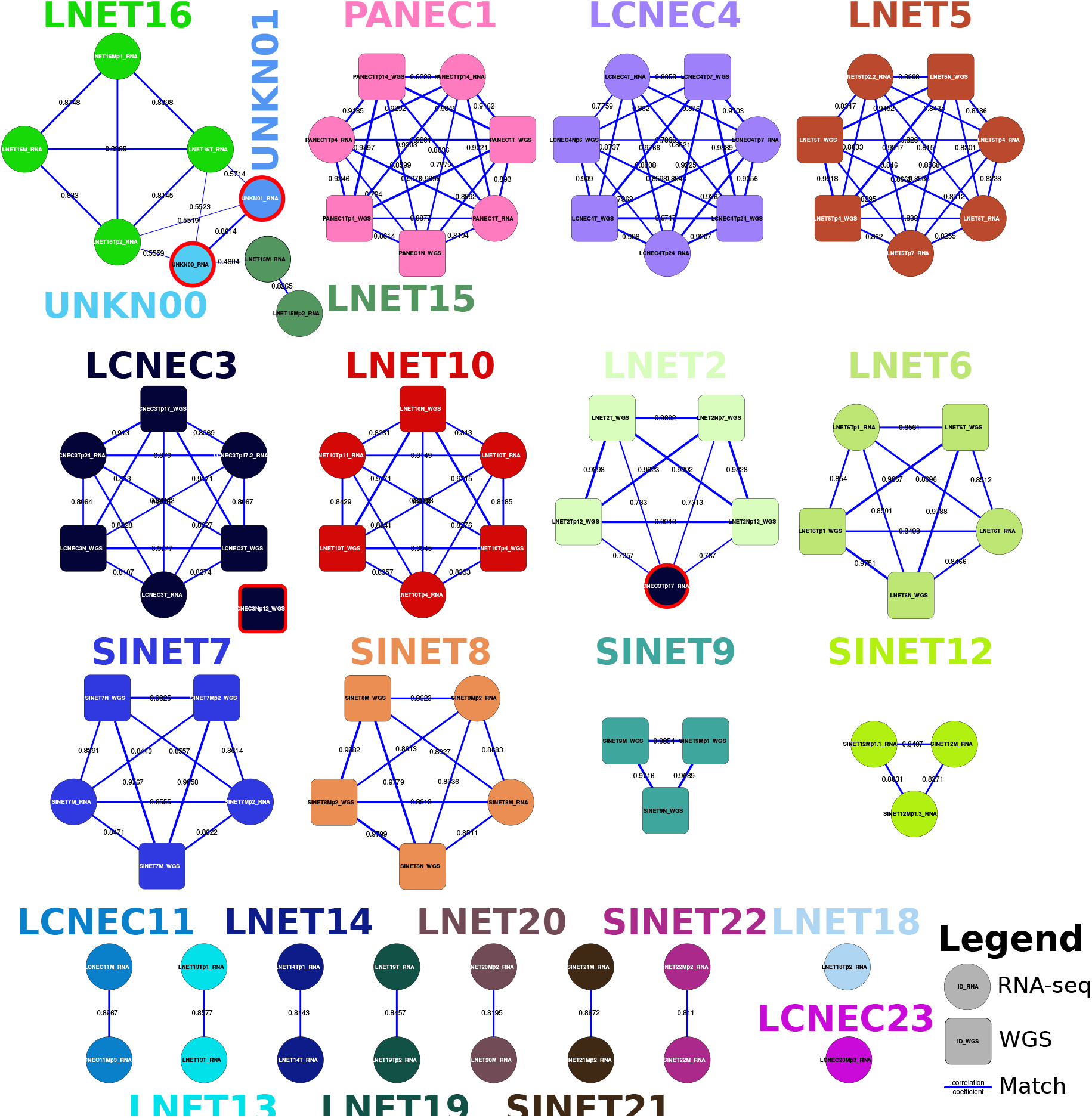
Network of matches between WGS and RNA-seq samples, computed with software NGSCheckmate. Numbers on the edges and edge thickness correspond to the Pearson correlation coeicient *r* between allelic fractions for the germline SNP panel; colors: experiments (see Table 1); squares: WGS, circles: RNA-seq, red contour: mismatches.

### Sex validation

We validated the sex reported in the clinical data using the multi-omic data. For the WGS data, we used the proportion of reads aligned to the sex chromosomes to assess whether samples clustered by sex (Fig. 5A). We found that all samples clustered by sex except for the normal of experiment LCNEC3 (sample LCNEC3Np12) which clustered with females despite other samples from the experiment clearly clustering with males. This further supports the sample matching reports that suggest that this sample does not match the rest of the experiment. For the RNA-seq data, we compared the total expression level on the sex chromosomes, using the variance-stabilised read counts as a quantification of gene expression (vst function from R package DESeq2 v1.26.0 [34]) (Fig. 5B). We find that samples from the same sex cluster together for all experiments, suggesting concordance with the clinical data.

**Figure 5.**
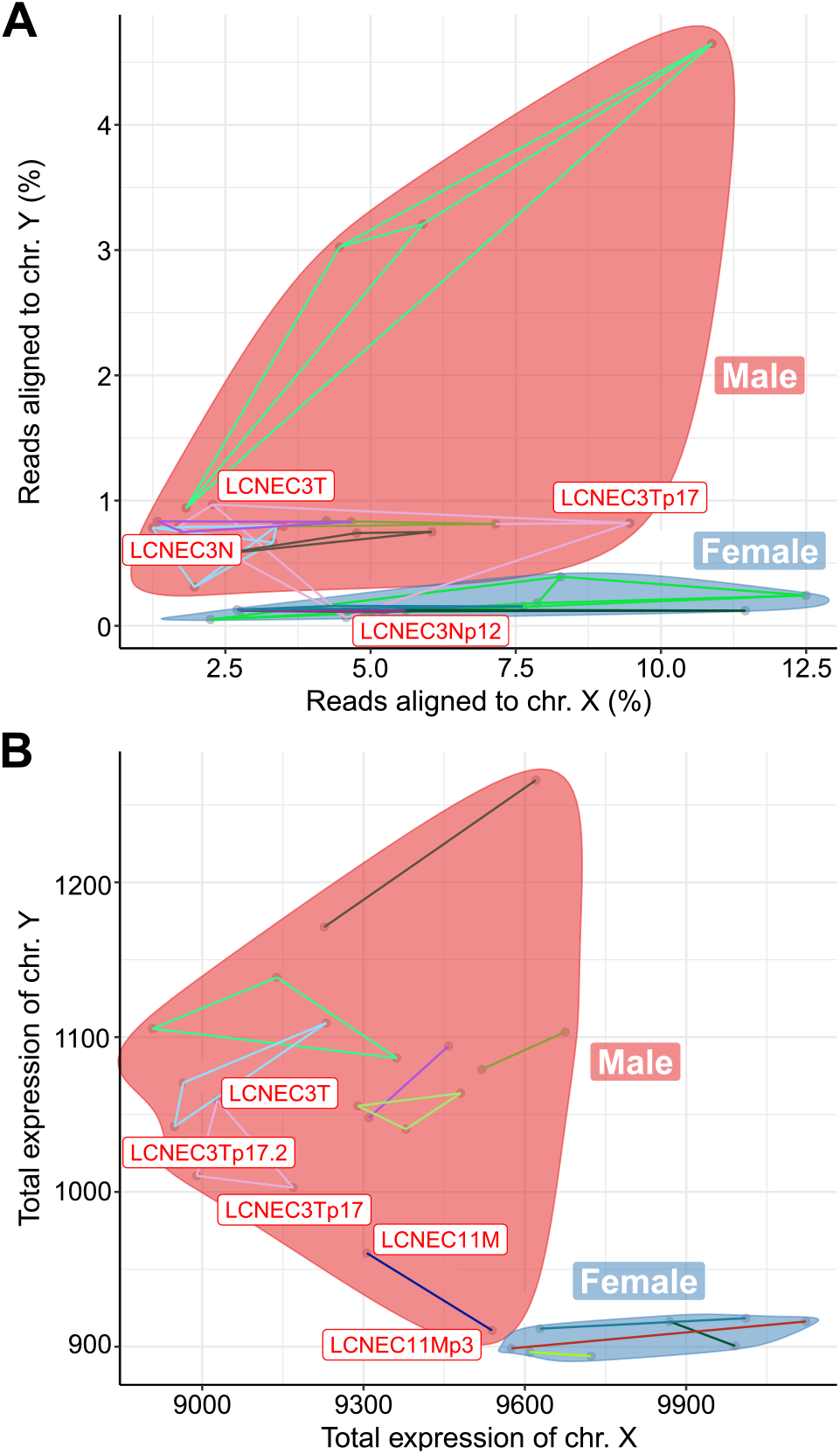
Validation of reported sex. (A) Percentage of reads aligned to chromosome X and Y in the whole-genome sequencing data. (B) Total gene expression in X and Y chromosome, in units of variance-stabilized read counts, computed from RNA-seq data. In all panels, samples from each sex are encircled (red:male, blue:female), excluding LCNEC3Np12, which we report as not matching the other samples from the LCNEC3 experiment.

### Small variant calls from RNA-seq

We classified small variants called from RNA-seq in 241 known neuroendocrine neoplasm driver genes (from Table S4 in [6]) as somatic or germline, using a random forest (RF) algorithm (R package randomForest v4.7-1.1 [36]; Fig. 6), using a similar approach as we recently did to classify mutations in tumor-only WGS [37]. After filtering out non-exonic, synonymous, and nonsynonymous mutations with a REVEL score [38] below 0.5, and mutations not in the list of 241 drivers, we were left with 2430 variants. Among them, 1174 variants were in samples with WGS data available and their somatic status was thus known.

**Figure 6.**
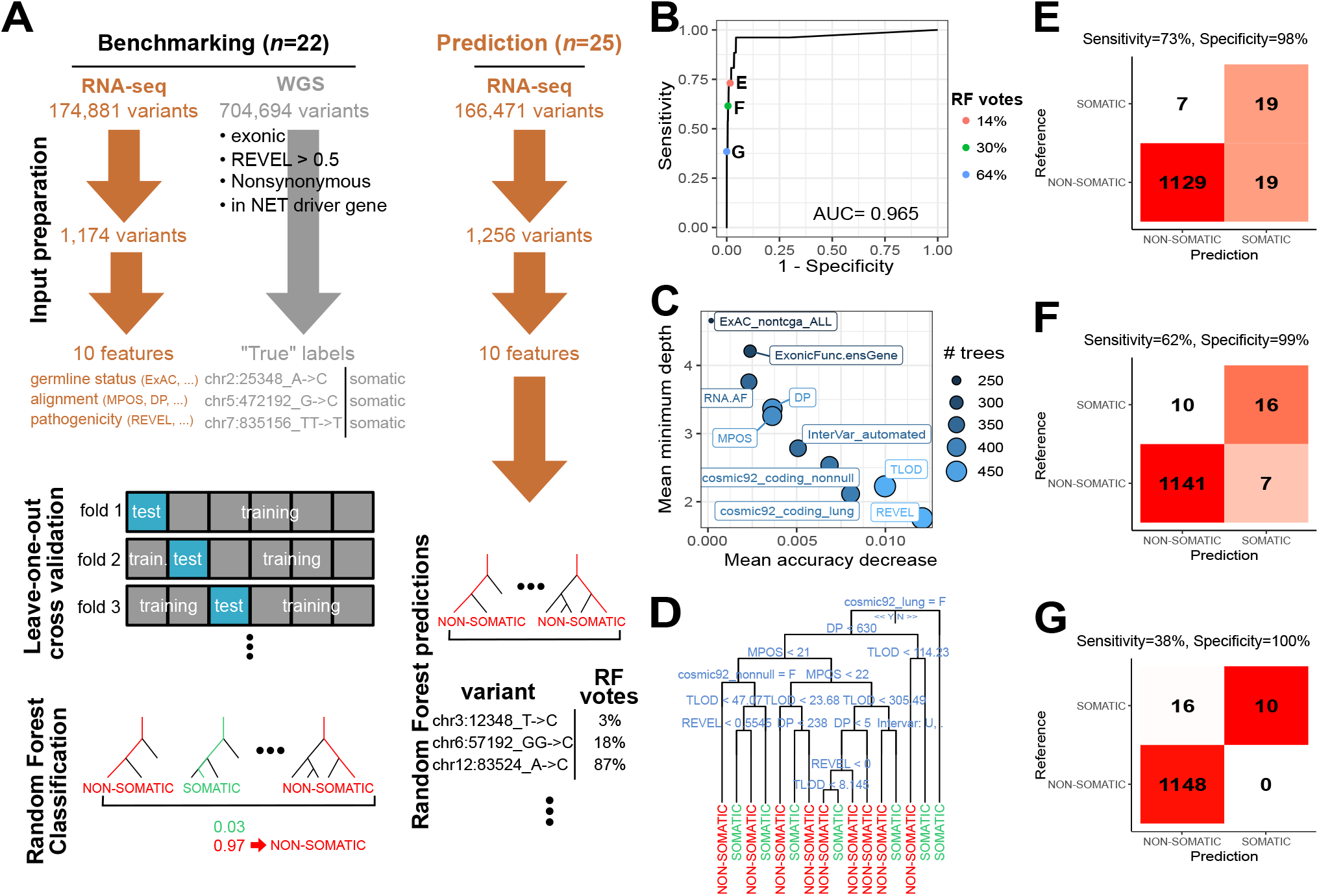
Random forest (RF) classification of variants as somatic or germline from RNA-seq data. (A) Schematic of the RF training, test, and prediction. B) Receiver operating characteristic (ROC) curve. C) Feature importance for classification accuracy. D) Representative tree of the RF. At each split, the split condition is written above, the left branch corresponds to a Yes and the right branch to a No. Final decision (SOMATIC or NON-SOMATIC) is represented by the leaves. (E)-(G) Confusion matrix for different levels of sensitivity and specificity. Reference: somatic status assessed from whole-genome sequencing data. Prediction: somatic status predicted from RNA-seq data using the RF algorithm.

We used 10 features in the RF model. One feature was directly informative about the potential germline status: the frequency of the allele in human populations from the ExAc database excluding cancers from the TCGA (feature *ExAC*_*nontcga*_*ALL*). Four features were informative about the alignment: the median distance from the end of the read (feature *MPOS*), the likelihood ratio score of variant existence (feature *TLOD*), the coverage at the position (feature *DP*), and the allelic fraction of the alternative allele (RNA.AF). Finally, the other features were informative about the pathogenicity of the variant: the REVEL score of pathogenicity (feature *REVEL*), the presence in the COSMIC 92 database (feature *cosmic*92_*coding*_*nonnull*), the presence in the COSMIC 92 database in a lung tumor (feature *cosmic*92_*coding*_*lung*), and the InterVar annotation (feature *InterVar*_*automated*; with levels “.”, “Uncertain_significance”, “likely_pathogenic”, and “Pathogenic”), and the exonic function of the variant (missense, nonsense, inframe or frameshift insertion, etc).

The RF algorithm was trained and tested on the 1174 variants with known status (1148 germline, 26 somatic) called in 22 samples from 8 experiments (Fig. 6A) We used leave-one-out cross-validation at the experiment level (8 folds), excluding all samples from one same experiment from the model fit at each iteration in order to avoid over-fitting due to the inclusion of variants from the same individual but different samples (e.g., LCNEC3T and LCNEC3Tp17) in the training and test sets. We used 5000 trees, and 3 features per split (the square root of the total number of features as recommended by default), and a minimal node size of 1. We estimated the performance of the model using the receiver operating characteristic (ROC) curve and its area under the curve (AUC, computed using the trapezoid rule), showing the sensitivity as a function of 1-specificity across different thresholds for the proportion of votes for the somatic class. We also computed the false discovery rate to get a sense of the proportion of variants classified as somatic that would actually be false positives. Once the RF model performance was assessed, we trained a RF model on the full 1174 variants and predicted the status of the remaining 1256 variants. See github repository https://github.com/IARCbioinfo/MS_panNEN_organoids for the complete R script.

We find that we can classify variants as somatic or germline with a balanced accuracy of 86%, with both specificity greater than 98% and sensitivity greater than 73% (AUC=0.965). Interestingly, although somatic variants are just a fraction of the calls (2%), the high sensitivities and specificities of our RF algorithm allowed to classify variants with false discovery rates below 50% while still preserving sensitivities above 60% (see Fig. 6B, E-G). We evaluated the importance of features for the classification both using the mean decrease in accuracy, which captures how much the model loses accuracy when the feature is excluded, and using the mean tree depth at which the feature was observed, with a low value meaning that the feature is used early in the decision trees and thus separates many variants[35, 39] (R package randomForestExplainer v0.10.1). The most important features for the classification were the REVEL score, the TLOD, and the cosmic annotation, while the frequency in the ExAC database was the least important, presumably because all these variants were very rare (Fig. 6C). Indeed, the most representative tree from the RF, computed using the reprtree R package v0.6 using the d2 distance metric between tree predictions[40], relied on these three variables, with all alterations present in a lung tumor from the COSMIC 92 database automatically classified as somatic (root of the tree), and TLOD and REVEL score being the most common features used for splitting (Fig. 6D).

## Re-use potential

We describe here some of the very first multi-omic datasets for patient-derived tumor organoids of pancreatic, small intestine (ileum), and pulmonary neuroendocrine neoplasms, in particular including the first lung neuroendocrine tumor organoids. Because such low grade tumors are diicult to cultivate in vitro, there is currently a lack of adequate experimental systems for these tumors, and we expect the biobank associated with the data presented here to be the basis for future experimental studies–either fundamental or treatment oriented–on neuroendocrine neoplasms across body sites. The multi-omic dataset we provide here constitutes the molecular fingerprints of these experimental models, and will be key to investigate oncogenic processes responsible for tumor initiation and progression, and to link drug responses to molecular features to design future personalized treatments.

To facilitate future studies, we used the exact same data processing as in our previous studies of neuroendocrine neoplasms [14, 15] and other rare cancers [41], in particular using rigorous RNA-seq expression quantification with containerized software and operating systems (see methods section). In addition, we provide all R scripts to analyze the data at https://github.com/IARCbioinfo/MS_panNEN_organoids.

## Conclusion

We have shown that our multi-omic dataset is of high quality and can be easily re-used. Given the rarity of neuroendocrine tumors from the lung, pancreas, and small intestine, past genomic studies each only reported data for a handful of samples, limiting the potential discoveries. For example, for lung NETs, 29 WGS and 39 RNAseq were reported in [42], 3 WGS and 20 RNA-seq in [14], and 30 RNA-seq in [43]; for small intestine NETs, for example, 81 RNA-seq with no WGS were reported in [44] and 7 RNA-seq in [45]). As a result, the primary tumors and metastasis sequencing data we report here (10 samples with WGS, 21 with RNA-seq) alone are very valuable, and should be combined with other datasets in future studies to provide enough power to discover informative molecular features for diagnosis, prognosis, and treatment. In addition, we report a unique multi-omic dataset generated from patient-derived tumor organoids, which will allow all researchers working on our biobank to test hypotheses regarding the molecular features associated with drug responses and thus advance research on personalized treatments for these understudied diseases.

## Availability of source code and requirements

- Project name: NEN organoids project, lungNENomics
- Project home pages: https://www.embl.org/groups/dayton/, http://rarecancersgenomics.com/lungnenomics/
- Operating system(s): Platform independent
- Programming language: Nextflow, R
- Other requirements: R packages *caret, randomForest*
- License: GNU GPL

## Availability of supporting data and materials

The data set supporting the results of this article is available in the European Genome-Phenome archive repository, study EGAS00001005752. The study consists of seven datasets: EGAD00001009988, with WGS CRAM files for 2 experiments, EGAD00001009989 with WGS CRAM files for 6 experiments, EGAD00001009990, with WGS CRAM files for 2 experiments, EGAD00001009991 with RNA-seq fastq files from 4 experiments, EGAD00001009992, with RNAseq fastq files for 15 experiments, EGAD00001009993, with RNA-seq fastq files for 2 experiments, and EGAD00001009994, with gene expression in multiple formats (R data, tab-separated text files) and multiple units (raw counts, TPM, FPKM) for 21 samples. Because of the sensitivity of the data and the patient consent, to get access to the data, please contact the data access committee of the Division of Biomedical Genetics from UMC Utrecht at. Once a data access agreement has been signed and access granted, data can be downloaded using the EGA python client (see detailed instructions at https://github.com/EGA-archive/ega-download-client, and video tutorial at https://embl-ebi.cloud.panopto.eu/Panopto/Pages/Viewer.aspx?id=be79bb93-1737-4f95-b80f-ab4300aa6f5a)

The multiQC report for WGS raw reads is available in Supplementary File S1; the multiQC report for RNA-seq raw reads is available in Supplementary File S2; the multiQC report for WGS alignments is available in Supplementary File S3; the multiQC report for RNA-seq alignments is available in Supplementary File S4.

## Abbreviations

bp: base pairs
LCNEC: large-cell neuroendocrine carcinoma
NEC: neuroendocrine carcinoma
NEN: neuroendocrine neoplasms
NET: neuroendocrine tumors
RNA-seq: RNA-sequencing
WGS: whole-genome sequencing
RF: random forest

## Ethical Approval

This study was approved by the medical ethical committee of of each respective hospital of the patients: Verenigde Commissies Mensgebonden Onderzoek of the St. Antonius Hospital Nieuwegein, Z-12.55; UMC Utrecht, METC 12-093 HUB-Cancer; NKI Institutional Review Board (IRB), M18ORG/CFMPB582; Maastricht University Medical Center, METC 2019-1061, and 2019-1039.

## Consent for publication

All patients signed informed consent forms for molecular analyses and to the publishing of the data.

## Author’s Contributions

TD designed the study and conducted the experiments. NA and MF designed the bioinformatic workflows. NA performed the data processing. NA and LM performed the analyses. CV formatted and deposited the data on EGA. MF, LFC, and TD supervised the analyses. NA, TD, AS-O, MF, and LFC wrote the manuscript. All authors reviewed and commented the paper.

## Acknowledgements

We thank the patients for participating to the study. We thank Utrecht Sequencing for RNA-sequencing services. The results shown here are in part based upon data generated by the Rare Cancers Genomics initiative (www.rarecancersgenomics.com).

