## Supplementary material for "Multi-omic dataset of patient-derived tumor organoids of neuroendocrine neoplasms": SupplementaryFileS2.html

NEN organoids: MultiQC Report


### Toggle navigation v1.7

### NEN organoids

Loading report..

- General Stats
- FastQC
  - Sequence Counts
  - Sequence Quality Histograms
  - Per Sequence Quality Scores
  - Per Base Sequence Content
  - Per Sequence GC Content
  - Per Base N Content
  - Sequence Length Distribution
  - Sequence Duplication Levels
  - Overrepresented sequences
  - Adapter Content

Toolbox

##### MultiQC Toolbox

###### Apply Highlight Samples

+

Regex mode off
help
 Clear

###### Apply Rename Samples

+

Click here for bulk input.

Paste two columns of a tab-delimited table here (eg. from Excel).

First column should be the old name, second column the new name.

Add

Regex mode off
help
 Clear

###### Apply Show / Hide Samples

Hide matching samples

Show only matching samples

+

Regex mode off
help
 Clear

###### Export Plots

- Images
- Data

px

px

Aspect ratio

PNG
JPEG
SVG

Plot scaling

X

Download the raw data used to create the plots in this report below:

Format:

Tab-separated
Comma-separated
JSON

Note that additional data was saved in `multiqc_pretrim_report_it_data` when this report was generated.

---

###### Choose Plots

 All
 None


---


   Download Plot Images

If you use plots from MultiQC in a publication or presentation, please cite:

> **MultiQC: Summarize analysis results for multiple tools and samples in a single report**  
> *Philip Ewels, Måns Magnusson, Sverker Lundin and Max Käller*  
> Bioinformatics (2016)  
> doi: 10.1093/bioinformatics/btw354  
> PMID: 27312411

###### Save Settings

You can save the toolbox settings for this report to the browser.

 Save


---

###### Load Settings

Choose a saved report profile from the dropdown box below:

[ select ]

Load
 Delete
 Set default
 Clear default

###### About MultiQC

This report was generated using MultiQC, version 1.7

You can see a YouTube video describing how to use MultiQC reports here:
https://youtu.be/qPbIlO\_KWN0

For more information about MultiQC, including other videos and
extensive documentation, please visit http://multiqc.info

You can report bugs, suggest improvements and find the source code for MultiQC on GitHub:
https://github.com/ewels/MultiQC

MultiQC is published in Bioinformatics:

> **MultiQC: Summarize analysis results for multiple tools and samples in a single report**  
> *Philip Ewels, Måns Magnusson, Sverker Lundin and Max Käller*  
> Bioinformatics (2016)  
> doi: 10.1093/bioinformatics/btw354  
> PMID: 27312411

### NEN organoids

A modular tool to aggregate results from bioinformatics analyses across many samples into a single report.

> RNA-seq Pre-trimming QC report

Contact E-mail
:  

Application Type
:   Multi-omic sequencing of lung, small intestine, and pancreas neuroendocrine neoplasm organoids

###### JavaScript Disabled

MultiQC reports use JavaScript for plots and toolbox functions. It looks like
you have JavaScript disabled in your web browser. Please note that many of the report
functions will not work as intended.

Loading report..

Report generated on 2020-07-15, 13:30 based on data in:
`/data/gcs/lungNENomics/work/organoids/RNAseq/processing/RNAseq-nf_organoids_13072020/QC/fastq`

Change sample names:
Sequencing Center IDs
IDs

---

×
don't show again

**Welcome!** Not sure where to start?  
Watch a tutorial video
  *(6:06)*

#### General Statistics

 Copy table

 Configure Columns

 Sort by highlight

 Plot
Showing ^140^/_140_ rows and ^4^/_5_ columns.

| Sample Name | % Dups | % GC | Length | % Failed | M Seqs |
| --- | --- | --- | --- | --- | --- |
| TD10B\_AHMVVKDRXX\_S8\_L001\_R1\_001 | 83.7% | 50% | 151 bp | 27% | 53.7 |
| TD10B\_AHMVVKDRXX\_S8\_L001\_R2\_001 | 74.2% | 49% | 151 bp | 18% | 53.7 |
| TD10B\_AHMVVKDRXX\_S8\_L002\_R1\_001 | 83.3% | 50% | 151 bp | 27% | 54.2 |
| TD10B\_AHMVVKDRXX\_S8\_L002\_R2\_001 | 73.8% | 49% | 151 bp | 18% | 54.2 |
| TD10C\_AHMVVKDRXX\_S9\_L001\_R1\_001 | 83.1% | 51% | 151 bp | 27% | 43.0 |
| TD10C\_AHMVVKDRXX\_S9\_L001\_R2\_001 | 67.5% | 49% | 151 bp | 36% | 43.0 |
| TD10C\_AHMVVKDRXX\_S9\_L002\_R1\_001 | 82.9% | 51% | 151 bp | 27% | 43.3 |
| TD10C\_AHMVVKDRXX\_S9\_L002\_R2\_001 | 67.1% | 49% | 151 bp | 36% | 43.3 |
| TD11B\_AHMVVKDRXX\_S6\_L001\_R1\_001 | 81.3% | 50% | 151 bp | 27% | 48.6 |
| TD11B\_AHMVVKDRXX\_S6\_L001\_R2\_001 | 72.5% | 49% | 151 bp | 27% | 48.6 |
| TD11B\_AHMVVKDRXX\_S6\_L002\_R1\_001 | 80.5% | 50% | 151 bp | 27% | 48.5 |
| TD11B\_AHMVVKDRXX\_S6\_L002\_R2\_001 | 71.6% | 49% | 151 bp | 27% | 48.5 |
| TD11C\_AHMVVKDRXX\_S7\_L001\_R1\_001 | 79.5% | 51% | 151 bp | 27% | 50.7 |
| TD11C\_AHMVVKDRXX\_S7\_L001\_R2\_001 | 67.7% | 51% | 151 bp | 18% | 50.7 |
| TD11C\_AHMVVKDRXX\_S7\_L002\_R1\_001 | 79.1% | 51% | 151 bp | 27% | 51.3 |
| TD11C\_AHMVVKDRXX\_S7\_L002\_R2\_001 | 67.4% | 51% | 151 bp | 18% | 51.3 |
| TD12B\_AHMVVKDRXX\_S15\_L001\_R1\_001 | 77.8% | 51% | 151 bp | 27% | 37.3 |
| TD12B\_AHMVVKDRXX\_S15\_L001\_R2\_001 | 65.0% | 50% | 151 bp | 18% | 37.3 |
| TD12B\_AHMVVKDRXX\_S15\_L002\_R1\_001 | 77.4% | 51% | 151 bp | 27% | 37.6 |
| TD12B\_AHMVVKDRXX\_S15\_L002\_R2\_001 | 64.5% | 50% | 151 bp | 18% | 37.6 |
| TD12C\_AHMVVKDRXX\_S16\_L001\_R1\_001 | 86.1% | 50% | 151 bp | 27% | 38.6 |
| TD12C\_AHMVVKDRXX\_S16\_L001\_R2\_001 | 74.5% | 49% | 151 bp | 27% | 38.6 |
| TD12C\_AHMVVKDRXX\_S16\_L002\_R1\_001 | 85.7% | 51% | 151 bp | 27% | 38.8 |
| TD12C\_AHMVVKDRXX\_S16\_L002\_R2\_001 | 74.0% | 49% | 151 bp | 27% | 38.8 |
| TD12D\_AHMVVKDRXX\_S17\_L001\_R1\_001 | 77.2% | 51% | 151 bp | 27% | 47.8 |
| TD12D\_AHMVVKDRXX\_S17\_L001\_R2\_001 | 66.8% | 51% | 151 bp | 18% | 47.8 |
| TD12D\_AHMVVKDRXX\_S17\_L002\_R1\_001 | 76.6% | 51% | 151 bp | 27% | 48.4 |
| TD12D\_AHMVVKDRXX\_S17\_L002\_R2\_001 | 66.1% | 51% | 151 bp | 18% | 48.4 |
| TD13B\_AHMVVKDRXX\_S18\_L001\_R1\_001 | 80.7% | 50% | 151 bp | 27% | 42.6 |
| TD13B\_AHMVVKDRXX\_S18\_L001\_R2\_001 | 70.0% | 49% | 151 bp | 18% | 42.6 |
| TD13B\_AHMVVKDRXX\_S18\_L002\_R1\_001 | 80.1% | 51% | 151 bp | 27% | 43.0 |
| TD13B\_AHMVVKDRXX\_S18\_L002\_R2\_001 | 69.5% | 50% | 151 bp | 18% | 43.0 |
| TD13C\_AHMVVKDRXX\_S19\_L001\_R1\_001 | 80.3% | 50% | 151 bp | 27% | 53.2 |
| TD13C\_AHMVVKDRXX\_S19\_L001\_R2\_001 | 70.2% | 50% | 151 bp | 27% | 53.2 |
| TD13C\_AHMVVKDRXX\_S19\_L002\_R1\_001 | 79.8% | 51% | 151 bp | 27% | 53.6 |
| TD13C\_AHMVVKDRXX\_S19\_L002\_R2\_001 | 69.5% | 50% | 151 bp | 27% | 53.6 |
| TD14B\_AHMVVKDRXX\_S20\_L001\_R1\_001 | 81.0% | 51% | 151 bp | 27% | 43.5 |
| TD14B\_AHMVVKDRXX\_S20\_L001\_R2\_001 | 68.6% | 51% | 151 bp | 18% | 43.5 |
| TD14B\_AHMVVKDRXX\_S20\_L002\_R1\_001 | 80.5% | 52% | 151 bp | 27% | 43.9 |
| TD14B\_AHMVVKDRXX\_S20\_L002\_R2\_001 | 68.3% | 51% | 151 bp | 18% | 43.9 |
| TD14C\_AHMVVKDRXX\_S21\_L001\_R1\_001 | 87.3% | 52% | 151 bp | 36% | 36.8 |
| TD14C\_AHMVVKDRXX\_S21\_L001\_R2\_001 | 76.2% | 51% | 151 bp | 18% | 36.8 |
| TD14C\_AHMVVKDRXX\_S21\_L002\_R1\_001 | 87.0% | 52% | 151 bp | 36% | 37.1 |
| TD14C\_AHMVVKDRXX\_S21\_L002\_R2\_001 | 75.9% | 51% | 151 bp | 18% | 37.1 |
| TD1A\_HVYYHBGXB\_S6\_L001\_R1\_001 | 41.5% | 49% | 139 bp | 9% | 8.2 |
| TD1A\_HVYYHBGXB\_S6\_L001\_R2\_001 | 29.4% | 50% | 150 bp | 9% | 8.2 |
| TD1A\_HVYYHBGXB\_S6\_L002\_R1\_001 | 39.4% | 49% | 139 bp | 9% | 8.2 |
| TD1A\_HVYYHBGXB\_S6\_L002\_R2\_001 | 26.6% | 50% | 150 bp | 9% | 8.2 |
| TD1A\_HVYYHBGXB\_S6\_L003\_R1\_001 | 39.5% | 49% | 139 bp | 9% | 7.7 |
| TD1A\_HVYYHBGXB\_S6\_L003\_R2\_001 | 26.3% | 50% | 150 bp | 9% | 7.7 |
| TD1A\_HVYYHBGXB\_S6\_L004\_R1\_001 | 39.5% | 49% | 139 bp | 9% | 7.6 |
| TD1A\_HVYYHBGXB\_S6\_L004\_R2\_001 | 26.8% | 50% | 150 bp | 9% | 7.6 |
| TD1B\_HVYYHBGXB\_S7\_L001\_R1\_001 | 53.9% | 48% | 137 bp | 18% | 16.4 |
| TD1B\_HVYYHBGXB\_S7\_L001\_R2\_001 | 41.3% | 49% | 150 bp | 9% | 16.4 |
| TD1B\_HVYYHBGXB\_S7\_L002\_R1\_001 | 51.3% | 48% | 138 bp | 18% | 16.2 |
| TD1B\_HVYYHBGXB\_S7\_L002\_R2\_001 | 37.7% | 49% | 150 bp | 9% | 16.2 |
| TD1B\_HVYYHBGXB\_S7\_L003\_R1\_001 | 51.0% | 48% | 138 bp | 18% | 15.2 |
| TD1B\_HVYYHBGXB\_S7\_L003\_R2\_001 | 36.6% | 49% | 150 bp | 9% | 15.2 |
| TD1B\_HVYYHBGXB\_S7\_L004\_R1\_001 | 50.8% | 48% | 138 bp | 18% | 15.2 |
| TD1B\_HVYYHBGXB\_S7\_L004\_R2\_001 | 37.0% | 49% | 150 bp | 9% | 15.2 |
| TD1C\_HVYYHBGXB\_S1\_L001\_R1\_001 | 50.7% | 49% | 139 bp | 9% | 14.2 |
| TD1C\_HVYYHBGXB\_S1\_L001\_R2\_001 | 37.8% | 50% | 150 bp | 9% | 14.2 |
| TD1C\_HVYYHBGXB\_S1\_L002\_R1\_001 | 47.8% | 49% | 139 bp | 0% | 14.2 |
| TD1C\_HVYYHBGXB\_S1\_L002\_R2\_001 | 35.2% | 49% | 150 bp | 9% | 14.2 |
| TD1C\_HVYYHBGXB\_S1\_L003\_R1\_001 | 47.6% | 49% | 140 bp | 0% | 13.4 |
| TD1C\_HVYYHBGXB\_S1\_L003\_R2\_001 | 33.7% | 49% | 150 bp | 9% | 13.4 |
| TD1C\_HVYYHBGXB\_S1\_L004\_R1\_001 | 47.7% | 49% | 140 bp | 0% | 13.4 |
| TD1C\_HVYYHBGXB\_S1\_L004\_R2\_001 | 34.4% | 49% | 150 bp | 9% | 13.4 |
| TD3A\_HVYYHBGXB\_S2\_L001\_R1\_001 | 47.9% | 49% | 142 bp | 9% | 11.6 |
| TD3A\_HVYYHBGXB\_S2\_L001\_R2\_001 | 35.8% | 50% | 150 bp | 9% | 11.6 |
| TD3A\_HVYYHBGXB\_S2\_L002\_R1\_001 | 45.5% | 49% | 142 bp | 9% | 11.6 |
| TD3A\_HVYYHBGXB\_S2\_L002\_R2\_001 | 32.4% | 50% | 150 bp | 9% | 11.6 |
| TD3A\_HVYYHBGXB\_S2\_L003\_R1\_001 | 45.5% | 49% | 142 bp | 9% | 10.9 |
| TD3A\_HVYYHBGXB\_S2\_L003\_R2\_001 | 32.4% | 50% | 150 bp | 9% | 10.9 |
| TD3A\_HVYYHBGXB\_S2\_L004\_R1\_001 | 45.4% | 49% | 142 bp | 9% | 10.9 |
| TD3A\_HVYYHBGXB\_S2\_L004\_R2\_001 | 33.2% | 50% | 150 bp | 9% | 10.9 |
| TD3B\_HVYYHBGXB\_S3\_L001\_R1\_001 | 65.9% | 50% | 138 bp | 18% | 14.9 |
| TD3B\_HVYYHBGXB\_S3\_L001\_R2\_001 | 53.1% | 50% | 150 bp | 18% | 14.9 |
| TD3B\_HVYYHBGXB\_S3\_L002\_R1\_001 | 62.9% | 50% | 138 bp | 18% | 14.9 |
| TD3B\_HVYYHBGXB\_S3\_L002\_R2\_001 | 49.2% | 50% | 150 bp | 9% | 14.9 |
| TD3B\_HVYYHBGXB\_S3\_L003\_R1\_001 | 62.8% | 50% | 139 bp | 18% | 14.0 |
| TD3B\_HVYYHBGXB\_S3\_L003\_R2\_001 | 48.1% | 50% | 150 bp | 9% | 14.0 |
| TD3B\_HVYYHBGXB\_S3\_L004\_R1\_001 | 63.1% | 50% | 139 bp | 18% | 14.0 |
| TD3B\_HVYYHBGXB\_S3\_L004\_R2\_001 | 49.1% | 50% | 150 bp | 9% | 14.0 |
| TD3E\_AHMVVKDRXX\_S1\_L001\_R1\_001 | 80.6% | 52% | 151 bp | 27% | 52.1 |
| TD3E\_AHMVVKDRXX\_S1\_L001\_R2\_001 | 70.1% | 51% | 151 bp | 27% | 52.1 |
| TD3E\_AHMVVKDRXX\_S1\_L002\_R1\_001 | 80.4% | 52% | 151 bp | 27% | 52.7 |
| TD3E\_AHMVVKDRXX\_S1\_L002\_R2\_001 | 69.9% | 51% | 151 bp | 27% | 52.7 |
| TD4A\_HVYYHBGXB\_S5\_L001\_R1\_001 | 51.7% | 49% | 140 bp | 18% | 13.7 |
| TD4A\_HVYYHBGXB\_S5\_L001\_R2\_001 | 38.9% | 50% | 150 bp | 9% | 13.7 |
| TD4A\_HVYYHBGXB\_S5\_L002\_R1\_001 | 49.1% | 49% | 140 bp | 9% | 13.7 |
| TD4A\_HVYYHBGXB\_S5\_L002\_R2\_001 | 35.8% | 50% | 150 bp | 9% | 13.7 |
| TD4A\_HVYYHBGXB\_S5\_L003\_R1\_001 | 49.0% | 49% | 140 bp | 9% | 12.9 |
| TD4A\_HVYYHBGXB\_S5\_L003\_R2\_001 | 35.2% | 50% | 150 bp | 9% | 12.9 |
| TD4A\_HVYYHBGXB\_S5\_L004\_R1\_001 | 49.2% | 49% | 140 bp | 9% | 12.9 |
| TD4A\_HVYYHBGXB\_S5\_L004\_R2\_001 | 36.1% | 50% | 150 bp | 9% | 12.9 |
| TD4D\_AHMVVKDRXX\_S14\_L001\_R1\_001 | 80.4% | 50% | 151 bp | 27% | 46.3 |
| TD4D\_AHMVVKDRXX\_S14\_L001\_R2\_001 | 69.4% | 49% | 151 bp | 27% | 46.3 |
| TD4D\_AHMVVKDRXX\_S14\_L002\_R1\_001 | 80.0% | 50% | 151 bp | 27% | 46.1 |
| TD4D\_AHMVVKDRXX\_S14\_L002\_R2\_001 | 68.6% | 49% | 151 bp | 27% | 46.1 |
| TD4\_HVYYHBGXB\_S4\_L001\_R1\_001 | 67.8% | 48% | 140 bp | 18% | 15.0 |
| TD4\_HVYYHBGXB\_S4\_L001\_R2\_001 | 54.3% | 49% | 150 bp | 18% | 15.0 |
| TD4\_HVYYHBGXB\_S4\_L002\_R1\_001 | 65.2% | 48% | 140 bp | 18% | 14.9 |
| TD4\_HVYYHBGXB\_S4\_L002\_R2\_001 | 50.9% | 49% | 150 bp | 18% | 14.9 |
| TD4\_HVYYHBGXB\_S4\_L003\_R1\_001 | 65.2% | 48% | 140 bp | 18% | 14.0 |
| TD4\_HVYYHBGXB\_S4\_L003\_R2\_001 | 49.9% | 49% | 150 bp | 9% | 14.0 |
| TD4\_HVYYHBGXB\_S4\_L004\_R1\_001 | 65.4% | 48% | 140 bp | 18% | 14.0 |
| TD4\_HVYYHBGXB\_S4\_L004\_R2\_001 | 50.8% | 49% | 150 bp | 18% | 14.0 |
| TD5B\_AHMVVKDRXX\_S2\_L001\_R1\_001 | 83.1% | 51% | 151 bp | 27% | 56.0 |
| TD5B\_AHMVVKDRXX\_S2\_L001\_R2\_001 | 75.1% | 50% | 151 bp | 18% | 56.0 |
| TD5B\_AHMVVKDRXX\_S2\_L002\_R1\_001 | 82.8% | 51% | 151 bp | 27% | 56.9 |
| TD5B\_AHMVVKDRXX\_S2\_L002\_R2\_001 | 74.7% | 51% | 151 bp | 18% | 56.9 |
| TD5C\_AHMVVKDRXX\_S3\_L001\_R1\_001 | 79.5% | 51% | 151 bp | 27% | 50.7 |
| TD5C\_AHMVVKDRXX\_S3\_L001\_R2\_001 | 70.2% | 50% | 151 bp | 18% | 50.7 |
| TD5C\_AHMVVKDRXX\_S3\_L002\_R1\_001 | 79.4% | 51% | 151 bp | 27% | 51.3 |
| TD5C\_AHMVVKDRXX\_S3\_L002\_R2\_001 | 70.0% | 51% | 151 bp | 18% | 51.3 |
| TD6B\_AHMVVKDRXX\_S4\_L001\_R1\_001 | 78.9% | 51% | 151 bp | 27% | 54.6 |
| TD6B\_AHMVVKDRXX\_S4\_L001\_R2\_001 | 68.6% | 51% | 151 bp | 27% | 54.6 |
| TD6B\_AHMVVKDRXX\_S4\_L002\_R1\_001 | 78.4% | 51% | 151 bp | 27% | 55.1 |
| TD6B\_AHMVVKDRXX\_S4\_L002\_R2\_001 | 68.0% | 51% | 151 bp | 27% | 55.1 |
| TD6C\_AHMVVKDRXX\_S5\_L001\_R1\_001 | 78.6% | 51% | 151 bp | 27% | 53.3 |
| TD6C\_AHMVVKDRXX\_S5\_L001\_R2\_001 | 68.9% | 51% | 151 bp | 18% | 53.3 |
| TD6C\_AHMVVKDRXX\_S5\_L002\_R1\_001 | 78.6% | 51% | 151 bp | 27% | 54.1 |
| TD6C\_AHMVVKDRXX\_S5\_L002\_R2\_001 | 68.4% | 51% | 151 bp | 18% | 54.1 |
| TD7B\_AHMVVKDRXX\_S10\_L001\_R1\_001 | 77.6% | 51% | 151 bp | 27% | 44.5 |
| TD7B\_AHMVVKDRXX\_S10\_L001\_R2\_001 | 66.8% | 51% | 151 bp | 18% | 44.5 |
| TD7B\_AHMVVKDRXX\_S10\_L002\_R1\_001 | 77.3% | 51% | 151 bp | 27% | 45.0 |
| TD7B\_AHMVVKDRXX\_S10\_L002\_R2\_001 | 66.5% | 51% | 151 bp | 18% | 45.0 |
| TD7C\_AHMVVKDRXX\_S11\_L001\_R1\_001 | 79.6% | 52% | 151 bp | 27% | 33.7 |
| TD7C\_AHMVVKDRXX\_S11\_L001\_R2\_001 | 64.5% | 51% | 151 bp | 36% | 33.7 |
| TD7C\_AHMVVKDRXX\_S11\_L002\_R1\_001 | 79.3% | 52% | 151 bp | 27% | 34.1 |
| TD7C\_AHMVVKDRXX\_S11\_L002\_R2\_001 | 64.2% | 51% | 151 bp | 36% | 34.1 |
| TD8B\_AHMVVKDRXX\_S12\_L001\_R1\_001 | 78.3% | 51% | 151 bp | 27% | 46.6 |
| TD8B\_AHMVVKDRXX\_S12\_L001\_R2\_001 | 68.8% | 52% | 151 bp | 27% | 46.6 |
| TD8B\_AHMVVKDRXX\_S12\_L002\_R1\_001 | 77.6% | 51% | 151 bp | 27% | 46.9 |
| TD8B\_AHMVVKDRXX\_S12\_L002\_R2\_001 | 68.1% | 52% | 151 bp | 27% | 46.9 |
| TD8C\_AHMVVKDRXX\_S13\_L001\_R1\_001 | 78.5% | 52% | 151 bp | 27% | 44.4 |
| TD8C\_AHMVVKDRXX\_S13\_L001\_R2\_001 | 68.8% | 51% | 151 bp | 18% | 44.4 |
| TD8C\_AHMVVKDRXX\_S13\_L002\_R1\_001 | 78.2% | 52% | 151 bp | 27% | 44.9 |
| TD8C\_AHMVVKDRXX\_S13\_L002\_R2\_001 | 68.4% | 51% | 151 bp | 18% | 44.9 |

loading..

---

##### Sequence Length Distribution

The distribution of fragment sizes (read lengths) found.
See the FastQC help

loading..

---

##### Sequence Duplication Levels Help

The relative level of duplication found for every sequence.

loading..

**MultiQC v1.7** 
- Written by Phil Ewels,
available on GitHub.

This report uses HighCharts,
jQuery,
jQuery UI,
Bootstrap,
FileSaver.js and
clipboard.js.

×

##### Plot Table Data

Select Column

Select Column

Please select two table columns.

Close

×

##### Regex Help

Toolbox search strings can behave as regular expressions (regexes). Click a button below to see an example of it in action. Try modifying them yourself in the text box.

`^` (start of string)
`$` (end of string)
`[]` (character choice)
`\d` (shorthand for `[0-9]`)
`\w` (shorthand for `[0-9a-zA-Z_]`)
`.` (any character)
`\.` (literal full stop)
`()` `|` (group / separator)
`*` (prev char 0 or more)
`+` (prev char 1 or more)
`?` (prev char 0 or 1)
`{}` (char num times)
`{,}` (count range)

```
samp_1
samp_1_edited
samp_2
samp_2_edited
samp_3
samp_3_edited
prepended_samp_1
tmp_samp_1_edited
tmpp_samp_1_edited
tmppp_samp_1_edited
#samp_1_edited.tmp
samp_11
samp_11111
```

See regex101.com for a more heavy duty testing suite.

Close
