## Supplementary material for "Multi-omic dataset of patient-derived tumor organoids of neuroendocrine neoplasms": SupplementaryFileS3.html

NEN organoids: MultiQC Report


### Toggle navigation v1.9

### NEN organoids

Loading report..

- General Stats
- QualiMap
  - Coverage histogram
  - Cumulative genome coverage
  - Insert size histogram
  - GC content distribution
- GATK
- Samtools

Toolbox

##### MultiQC Toolbox

###### Apply Highlight Samples

+

Regex mode off
help
 Clear

###### Apply Rename Samples

+

Click here for bulk input.

Paste two columns of a tab-delimited table here (eg. from Excel).

First column should be the old name, second column the new name.

Add

Regex mode off
help
 Clear

###### Apply Show / Hide Samples

Hide matching samples

Show only matching samples

+

Regex mode off
help
 Clear

###### Export Plots

- Images
- Data

px

px

Aspect ratio

PNG
JPEG
SVG

Plot scaling

X

Download the raw data used to create the plots in this report below:

Format:

Tab-separated
Comma-separated
JSON

Note that additional data was saved in `multiqc_qualimap_flagstat_BQSR_report_data` when this report was generated.

---

###### Choose Plots

 All
 None


---


   Download Plot Images

If you use plots from MultiQC in a publication or presentation, please cite:

> **MultiQC: Summarize analysis results for multiple tools and samples in a single report**  
> *Philip Ewels, Måns Magnusson, Sverker Lundin and Max Käller*  
> Bioinformatics (2016)  
> doi: 10.1093/bioinformatics/btw354  
> PMID: 27312411

###### Save Settings

You can save the toolbox settings for this report to the browser.

 Save


---

###### Load Settings

Choose a saved report profile from the dropdown box below:

[ select ]

Load
 Delete
 Set default
 Clear default

###### About MultiQC

This report was generated using MultiQC, version 1.9

You can see a YouTube video describing how to use MultiQC reports here:
https://youtu.be/qPbIlO\_KWN0

For more information about MultiQC, including other videos and
extensive documentation, please visit http://multiqc.info

You can report bugs, suggest improvements and find the source code for MultiQC on GitHub:
https://github.com/ewels/MultiQC

MultiQC is published in Bioinformatics:

> **MultiQC: Summarize analysis results for multiple tools and samples in a single report**  
> *Philip Ewels, Måns Magnusson, Sverker Lundin and Max Käller*  
> Bioinformatics (2016)  
> doi: 10.1093/bioinformatics/btw354  
> PMID: 27312411

### NEN organoids

A modular tool to aggregate results from bioinformatics analyses across many samples into a single report.

> WGS/WES final QC report

Contact E-mail
:  

Application Type
:   Multi-omic sequencing of lung, small intestine, and pancreas neuroendocrine neoplasm organoids

###### JavaScript Disabled

MultiQC reports use JavaScript for plots and toolbox functions. It looks like
you have JavaScript disabled in your web browser. Please note that many of the report
functions will not work as intended.

Loading report..

Report
generated on 2020-09-02, 14:29
based on data in:
`/data/gcs/lungNENomics/work/organoids/WGS/alignment/QC`

Change sample names:
Sequencing Center IDs
IDs

---

×
don't show again

**Welcome!** Not sure where to start?  
Watch a tutorial video
  *(6:06)*

#### General Statistics

Showing 522 samples.

loading..

#### QualiMap

QualiMap is a platform-independent application to facilitate the quality control of alignment sequencing data and its derivatives like feature counts.

##### Coverage histogram Help

Distribution of the number of locations in the reference genome with a given depth of coverage.

For a set of DNA or RNA reads mapped to a reference sequence, such as a genome
or transcriptome, the depth of coverage at a given base position is the number
of high-quality reads that map to the reference at that position
(Sims et al. 2014).

Bases of a reference sequence (y-axis) are groupped by their depth of coverage
(*0×, 1×, …, N×*) (x-axis). This plot shows
the frequency of coverage depths relative to the reference sequence for each
read dataset, which provides an indirect measure of the level and variation of
coverage depth in the corresponding sequenced sample.

If reads are randomly distributed across the reference sequence, this plot
should resemble a Poisson distribution (Lander & Waterman 1988), with a peak indicating approximate
depth of coverage, and more uniform coverage depth being reflected in a narrower
spread. The optimal level of coverage depth depends on the aims of the
experiment, though it should at minimum be sufficiently high to adequately
address the biological question; greater uniformity of coverage is generally
desirable, because it increases breadth of coverage for a given depth of
coverage, allowing equivalent results to be achieved at a lower sequencing depth
(Sampson
et al. 2011; Sims
et al. 2014). However, it is difficult to achieve uniform coverage
depth in practice, due to biases introduced during sample preparation
(van
Dijk et al. 2014), sequencing (Ross et al. 2013) and read mapping
(Sims et al. 2014).

This plot may include a small peak for regions of the reference sequence with
zero depth of coverage. Such regions may be absent from the given sample (due
to a deletion or structural rearrangement), present in the sample but not
successfully sequenced (due to bias in sequencing or preparation), or sequenced
but not successfully mapped to the reference (due to the choice of mapping
algorithm, the presence of repeat sequences, or mismatches caused by variants
or sequencing errors). Related factors cause most datasets to contain some
unmapped reads (Sims
et al. 2014).

loading..

---

##### Cumulative genome coverage Help

Percentage of the reference genome with at least the given depth of coverage.

For a set of DNA or RNA reads mapped to a reference sequence, such as a genome
or transcriptome, the depth of coverage at a given base position is the number
of high-quality reads that map to the reference at that position, while the
breadth of coverage is the fraction of the reference sequence to which reads
have been mapped with at least a given depth of coverage
(Sims et al. 2014).

Defining coverage breadth in terms of coverage depth is useful, because
sequencing experiments typically require a specific minimum depth of coverage
over the region of interest (Sims et al. 2014), so the extent of the reference sequence
that is amenable to analysis is constrained to lie within regions that have
sufficient depth. With inadequate sequencing breadth, it can be difficult to
distinguish the absence of a biological feature (such as a gene) from a lack
of data (Green 2007).

For increasing coverage depths (*1×, 2×, …, N×*),
coverage breadth is calculated as the percentage of the reference
sequence that is covered by at least that number of reads, then plots
coverage breadth (y-axis) against coverage depth (x-axis). This plot
shows the relationship between sequencing depth and breadth for each read
dataset, which can be used to gauge, for example, the likely effect of a
minimum depth filter on the fraction of a genome available for analysis.

loading..

---

##### Insert size histogram Help

Distribution of estimated insert sizes of mapped reads.

To overcome limitations in the length of DNA or RNA sequencing reads,
many sequencing instruments can produce two or more shorter reads from
one longer fragment in which the relative position of reads is
approximately known, such as paired-end or mate-pair reads
(Mardis 2013). Such techniques can extend the reach
of sequencing technology, allowing for more accurate placement of reads
(Reinert et al. 2015) and better resolution of repeat
regions (Reinert et al. 2015), as well as detection of
structural variation (Alkan et al. 2011) and chimeric transcripts
(Maher et al. 2009).

All these methods assume that the approximate size of an insert is known.
(Insert size can be defined as the length in bases of a sequenced DNA or
RNA fragment, excluding technical sequences such as adapters, which are
typically removed before alignment.) This plot allows for that assumption
to be assessed. With the set of mapped fragments for a given sample, QualiMap
groups the fragments by insert size, then plots the frequency of mapped
fragments (y-axis) over a range of insert sizes (x-axis). In an ideal case,
the distribution of fragment sizes for a sequencing library would culminate
in a single peak indicating average insert size, with a narrow spread
indicating highly consistent fragment lengths.

QualiMap calculates insert sizes as follows: for each fragment in which
every read mapped successfully to the same reference sequence, it
extracts the insert size from the `TLEN` field of the leftmost read
(see the Qualimap 2 documentation), where the `TLEN` (or
'observed Template LENgth') field contains 'the number of bases from the
leftmost mapped base to the rightmost mapped base'
(SAM
format specification). Note that because it is defined in terms of
alignment to a reference sequence, the value of the `TLEN` field may
differ from the insert size due to factors such as alignment clipping,
alignment errors, or structural variation or splicing in a gap between
reads from the same fragment.

loading..

---

##### GC content distribution Help

Each solid line represents the distribution of GC content of mapped reads for a given sample.

GC bias is the difference between the guanine-cytosine content
(GC-content) of a set of sequencing reads and the GC-content of the DNA
or RNA in the original sample. It is a well-known issue with sequencing
systems, and may be introduced by PCR amplification, among other factors
(Benjamini
& Speed 2012; Ross et al. 2013).

QualiMap calculates the GC-content of individual mapped reads, then
groups those reads by their GC-content (*1%, 2%, …, 100%*), and
plots the frequency of mapped reads (y-axis) at each level of GC-content
(x-axis). This plot shows the GC-content distribution of mapped reads
for each read dataset, which should ideally resemble that of the
original sample. It can be useful to display the GC-content distribution
of an appropriate reference sequence for comparison, and QualiMap has an
option to do this (see the Qualimap 2 documentation).

loading..

---

#### GATK

GATK is a toolkit offering a wide variety of tools with a primary focus on variant discovery and genotyping.

##### Observed Quality Scores Help

This plot shows the distribution of base quality scores in each sample before and after base quality score recalibration (BQSR). Applying BQSR should broaden the distribution of base quality scores.

For more information see the Broad's description of BQSR.

Pre-recalibration Count
Pre-recalibration Percent

loading..

---

#### Samtools

Samtools is a suite of programs for interacting with high-throughput sequencing data.

##### Samtools Flagstat

This module parses the output from `samtools flagstat`. All numbers in millions.

loading..

**MultiQC v1.9** 
- Written by Phil Ewels,
available on GitHub.

This report uses HighCharts,
jQuery,
jQuery UI,
Bootstrap,
FileSaver.js and
clipboard.js.

×

##### Plot Table Data

Select Column

Select Column

Please select two table columns.

Close

×

##### Regex Help

Toolbox search strings can behave as regular expressions (regexes). Click a button below to see an example of it in action. Try modifying them yourself in the text box.

`^` (start of string)
`$` (end of string)
`[]` (character choice)
`\d` (shorthand for `[0-9]`)
`\w` (shorthand for `[0-9a-zA-Z_]`)
`.` (any character)
`\.` (literal full stop)
`()` `|` (group / separator)
`*` (prev char 0 or more)
`+` (prev char 1 or more)
`?` (prev char 0 or 1)
`{}` (char num times)
`{,}` (count range)

```
samp_1
samp_1_edited
samp_2
samp_2_edited
samp_3
samp_3_edited
prepended_samp_1
tmp_samp_1_edited
tmpp_samp_1_edited
tmppp_samp_1_edited
#samp_1_edited.tmp
samp_11
samp_11111
```

See regex101.com for a more heavy duty testing suite.

Close
