## Supplementary material for "Multi-omic dataset of patient-derived tumor organoids of neuroendocrine neoplasms": SupplementaryFileS4.html

NEN organoids: MultiQC Report


### Toggle navigation v1.8

### NEN organoids

Loading report..

- General Stats
- RSeQC
  - Read Distribution
  - Junction Saturation
- HTSeq Count
- STAR
- Cutadapt
- FastQC
  - Sequence Counts
  - Sequence Quality Histograms
  - Per Sequence Quality Scores
  - Per Base Sequence Content
  - Per Sequence GC Content
  - Per Base N Content
  - Sequence Length Distribution
  - Sequence Duplication Levels
  - Overrepresented sequences
  - Adapter Content
  - Status Checks

Toolbox

##### MultiQC Toolbox

###### Apply Highlight Samples

+

Regex mode off
help
 Clear

###### Apply Rename Samples

+

Click here for bulk input.

Paste two columns of a tab-delimited table here (eg. from Excel).

First column should be the old name, second column the new name.

Add

Regex mode off
help
 Clear

###### Apply Show / Hide Samples

Hide matching samples

Show only matching samples

+

Regex mode off
help
 Clear

###### Export Plots

- Images
- Data

px

px

Aspect ratio

PNG
JPEG
SVG

Plot scaling

X

Download the raw data used to create the plots in this report below:

Format:

Tab-separated
Comma-separated
JSON

Note that additional data was saved in `multiqc_posttrim_report_interactive_data_1` when this report was generated.

---

###### Choose Plots

 All
 None


---


   Download Plot Images

If you use plots from MultiQC in a publication or presentation, please cite:

> **MultiQC: Summarize analysis results for multiple tools and samples in a single report**  
> *Philip Ewels, Måns Magnusson, Sverker Lundin and Max Käller*  
> Bioinformatics (2016)  
> doi: 10.1093/bioinformatics/btw354  
> PMID: 27312411

###### Save Settings

You can save the toolbox settings for this report to the browser.

 Save


---

###### Load Settings

Choose a saved report profile from the dropdown box below:

[ select ]

Load
 Delete
 Set default
 Clear default

###### About MultiQC

This report was generated using MultiQC, version 1.8

You can see a YouTube video describing how to use MultiQC reports here:
https://youtu.be/qPbIlO\_KWN0

For more information about MultiQC, including other videos and
extensive documentation, please visit http://multiqc.info

You can report bugs, suggest improvements and find the source code for MultiQC on GitHub:
https://github.com/ewels/MultiQC

MultiQC is published in Bioinformatics:

> **MultiQC: Summarize analysis results for multiple tools and samples in a single report**  
> *Philip Ewels, Måns Magnusson, Sverker Lundin and Max Käller*  
> Bioinformatics (2016)  
> doi: 10.1093/bioinformatics/btw354  
> PMID: 27312411

### NEN organoids

A modular tool to aggregate results from bioinformatics analyses across many samples into a single report.

> RNA-seq

Contact E-mail
:  

Application Type
:   Multi-omic sequencing of lung, small intestine, and pancreas neuroendocrine neoplasm organoids

###### JavaScript Disabled

MultiQC reports use JavaScript for plots and toolbox functions. It looks like
you have JavaScript disabled in your web browser. Please note that many of the report
functions will not work as intended.

Loading report..

Report generated on 2020-07-20, 17:36 based on data in:
`/scratch/alcalan/nextflow_work/20/6278f2c37a5209ff064553a2500982`

Change sample names:
Sequencing Center IDs
IDs

---

×
don't show again

**Welcome!** Not sure where to start?  
Watch a tutorial video
  *(6:06)*

#### General Statistics

 Copy table

 Configure Columns

 Sort by highlight

 Plot
Showing ^336^/_336_ rows and ^8^/_10_ columns.

| Sample Name | % Assigned | M Assigned | % Aligned | M Aligned | % Trimmed | % Dups | % GC | Length | % Failed | M Seqs |
| --- | --- | --- | --- | --- | --- | --- | --- | --- | --- | --- |
| LCNEC11MAHMVVKDRXX\_S7\_L001\_val\_1 |  |  |  |  |  | 79.0% | 51% | 135 bp | 18% | 50.1 |
| LCNEC11MAHMVVKDRXX\_S7\_L001\_val\_2 |  |  |  |  |  | 68.4% | 50% | 135 bp | 9% | 50.1 |
| LCNEC11MAHMVVKDRXX\_S7\_L002\_val\_1 |  |  |  |  |  | 78.6% | 51% | 135 bp | 18% | 50.7 |
| LCNEC11MAHMVVKDRXX\_S7\_L002\_val\_2 |  |  |  |  |  | 68.0% | 50% | 135 bp | 9% | 50.7 |
| LCNEC11M\_count.txt | 82.8% | 76.6 |  |  |  |  |  |  |  |  |
| LCNEC11Mp3AHMVVKDRXX\_S6\_L001\_val\_1 |  |  |  |  |  | 80.9% | 49% | 133 bp | 18% | 48.2 |
| LCNEC11Mp3AHMVVKDRXX\_S6\_L001\_val\_2 |  |  |  |  |  | 72.8% | 49% | 133 bp | 9% | 48.2 |
| LCNEC11Mp3AHMVVKDRXX\_S6\_L002\_val\_1 |  |  |  |  |  | 80.0% | 50% | 133 bp | 18% | 48.2 |
| LCNEC11Mp3AHMVVKDRXX\_S6\_L002\_val\_2 |  |  |  |  |  | 71.9% | 49% | 133 bp | 9% | 48.2 |
| LCNEC11Mp3\_count.txt | 78.1% | 68.3 |  |  |  |  |  |  |  |  |
| LCNEC3THVYYHBGXB\_S3\_L001\_val\_1 |  |  |  |  |  | 65.5% | 50% | 137 bp | 18% | 14.4 |
| LCNEC3THVYYHBGXB\_S3\_L001\_val\_2 |  |  |  |  |  | 54.8% | 50% | 136 bp | 18% | 14.4 |
| LCNEC3THVYYHBGXB\_S3\_L002\_val\_1 |  |  |  |  |  | 62.5% | 50% | 137 bp | 18% | 14.4 |
| LCNEC3THVYYHBGXB\_S3\_L002\_val\_2 |  |  |  |  |  | 50.6% | 50% | 136 bp | 18% | 14.4 |
| LCNEC3THVYYHBGXB\_S3\_L003\_val\_1 |  |  |  |  |  | 62.5% | 50% | 137 bp | 18% | 13.6 |
| LCNEC3THVYYHBGXB\_S3\_L003\_val\_2 |  |  |  |  |  | 49.7% | 50% | 136 bp | 9% | 13.6 |
| LCNEC3THVYYHBGXB\_S3\_L004\_val\_1 |  |  |  |  |  | 62.8% | 50% | 137 bp | 18% | 13.6 |
| LCNEC3THVYYHBGXB\_S3\_L004\_val\_2 |  |  |  |  |  | 50.7% | 50% | 136 bp | 18% | 13.6 |
| LCNEC3T\_count.txt | 73.2% | 36.1 |  |  |  |  |  |  |  |  |
| LCNEC3Tp17.2AHMVVKDRXX\_S1\_L001\_val\_1 |  |  |  |  |  | 80.0% | 51% | 131 bp | 18% | 51.5 |
| LCNEC3Tp17.2AHMVVKDRXX\_S1\_L001\_val\_2 |  |  |  |  |  | 70.8% | 51% | 131 bp | 9% | 51.5 |
| LCNEC3Tp17.2AHMVVKDRXX\_S1\_L002\_val\_1 |  |  |  |  |  | 79.7% | 52% | 131 bp | 18% | 52.1 |
| LCNEC3Tp17.2AHMVVKDRXX\_S1\_L002\_val\_2 |  |  |  |  |  | 70.5% | 51% | 131 bp | 9% | 52.1 |
| LCNEC3Tp17.2\_count.txt | 81.6% | 77.5 |  |  |  |  |  |  |  |  |
| LCNEC3Tp17HVYYHBGXB\_S2\_L001\_val\_1 |  |  |  |  |  | 47.5% | 49% | 140 bp | 9% | 11.2 |
| LCNEC3Tp17HVYYHBGXB\_S2\_L001\_val\_2 |  |  |  |  |  | 37.2% | 50% | 139 bp | 9% | 11.2 |
| LCNEC3Tp17HVYYHBGXB\_S2\_L002\_val\_1 |  |  |  |  |  | 45.2% | 49% | 141 bp | 9% | 11.2 |
| LCNEC3Tp17HVYYHBGXB\_S2\_L002\_val\_2 |  |  |  |  |  | 33.7% | 50% | 139 bp | 9% | 11.2 |
| LCNEC3Tp17HVYYHBGXB\_S2\_L003\_val\_1 |  |  |  |  |  | 45.1% | 49% | 140 bp | 9% | 10.6 |
| LCNEC3Tp17HVYYHBGXB\_S2\_L003\_val\_2 |  |  |  |  |  | 33.6% | 50% | 139 bp | 9% | 10.6 |
| LCNEC3Tp17HVYYHBGXB\_S2\_L004\_val\_1 |  |  |  |  |  | 45.0% | 49% | 140 bp | 9% | 10.5 |
| LCNEC3Tp17HVYYHBGXB\_S2\_L004\_val\_2 |  |  |  |  |  | 34.6% | 50% | 139 bp | 9% | 10.5 |
| LCNEC3Tp17\_count.txt | 79.4% | 30.4 |  |  |  |  |  |  |  |  |
| LCNEC4THVYYHBGXB\_S4\_L001\_val\_1 |  |  |  |  |  | 67.5% | 48% | 138 bp | 18% | 14.6 |
| LCNEC4THVYYHBGXB\_S4\_L001\_val\_2 |  |  |  |  |  | 55.6% | 48% | 138 bp | 18% | 14.6 |
| LCNEC4THVYYHBGXB\_S4\_L002\_val\_1 |  |  |  |  |  | 64.9% | 48% | 139 bp | 18% | 14.5 |
| LCNEC4THVYYHBGXB\_S4\_L002\_val\_2 |  |  |  |  |  | 51.9% | 48% | 138 bp | 18% | 14.5 |
| LCNEC4THVYYHBGXB\_S4\_L003\_val\_1 |  |  |  |  |  | 64.9% | 48% | 138 bp | 18% | 13.7 |
| LCNEC4THVYYHBGXB\_S4\_L003\_val\_2 |  |  |  |  |  | 51.3% | 48% | 138 bp | 18% | 13.7 |
| LCNEC4THVYYHBGXB\_S4\_L004\_val\_1 |  |  |  |  |  | 65.0% | 48% | 138 bp | 18% | 13.6 |
| LCNEC4THVYYHBGXB\_S4\_L004\_val\_2 |  |  |  |  |  | 52.2% | 48% | 138 bp | 18% | 13.6 |
| LCNEC4T\_count.txt | 76.1% | 38.2 |  |  |  |  |  |  |  |  |
| LCNEC4Tp24HVYYHBGXB\_S5\_L001\_val\_1 |  |  |  |  |  | 51.3% | 49% | 139 bp | 18% | 13.2 |
| LCNEC4Tp24HVYYHBGXB\_S5\_L001\_val\_2 |  |  |  |  |  | 40.6% | 49% | 137 bp | 9% | 13.2 |
| LCNEC4Tp24HVYYHBGXB\_S5\_L002\_val\_1 |  |  |  |  |  | 48.7% | 49% | 139 bp | 9% | 13.2 |
| LCNEC4Tp24HVYYHBGXB\_S5\_L002\_val\_2 |  |  |  |  |  | 37.2% | 49% | 138 bp | 9% | 13.2 |
| LCNEC4Tp24HVYYHBGXB\_S5\_L003\_val\_1 |  |  |  |  |  | 48.6% | 49% | 139 bp | 9% | 12.5 |
| LCNEC4Tp24HVYYHBGXB\_S5\_L003\_val\_2 |  |  |  |  |  | 36.8% | 49% | 137 bp | 9% | 12.5 |
| LCNEC4Tp24HVYYHBGXB\_S5\_L004\_val\_1 |  |  |  |  |  | 48.8% | 49% | 139 bp | 9% | 12.4 |
| LCNEC4Tp24HVYYHBGXB\_S5\_L004\_val\_2 |  |  |  |  |  | 37.7% | 49% | 137 bp | 9% | 12.4 |
| LCNEC4Tp24\_count.txt | 79.9% | 35.9 |  |  |  |  |  |  |  |  |
| LCNEC4Tp7AHMVVKDRXX\_S14\_L001\_val\_1 |  |  |  |  |  | 79.8% | 50% | 133 bp | 18% | 45.6 |
| LCNEC4Tp7AHMVVKDRXX\_S14\_L001\_val\_2 |  |  |  |  |  | 70.3% | 48% | 133 bp | 9% | 45.6 |
| LCNEC4Tp7AHMVVKDRXX\_S14\_L002\_val\_1 |  |  |  |  |  | 79.4% | 50% | 133 bp | 18% | 45.4 |
| LCNEC4Tp7AHMVVKDRXX\_S14\_L002\_val\_2 |  |  |  |  |  | 69.5% | 49% | 133 bp | 9% | 45.4 |
| LCNEC4Tp7\_count.txt | 80.6% | 66.2 |  |  |  |  |  |  |  |  |
| LNET10TAHMVVKDRXX\_S9\_L001\_val\_1 |  |  |  |  |  | 82.5% | 50% | 133 bp | 18% | 41.9 |
| LNET10TAHMVVKDRXX\_S9\_L001\_val\_2 |  |  |  |  |  | 69.5% | 49% | 133 bp | 27% | 41.9 |
| LNET10TAHMVVKDRXX\_S9\_L002\_val\_1 |  |  |  |  |  | 82.2% | 50% | 133 bp | 18% | 42.1 |
| LNET10TAHMVVKDRXX\_S9\_L002\_val\_2 |  |  |  |  |  | 69.2% | 49% | 133 bp | 27% | 42.1 |
| LNET10T\_count.txt | 79.2% | 55.7 |  |  |  |  |  |  |  |  |
| LNET10Tp4AHMVVKDRXX\_S8\_L001\_val\_1 |  |  |  |  |  | 83.2% | 49% | 134 bp | 18% | 53.1 |
| LNET10Tp4AHMVVKDRXX\_S8\_L001\_val\_2 |  |  |  |  |  | 75.0% | 48% | 134 bp | 9% | 53.1 |
| LNET10Tp4AHMVVKDRXX\_S8\_L002\_val\_1 |  |  |  |  |  | 82.8% | 49% | 134 bp | 18% | 53.6 |
| LNET10Tp4AHMVVKDRXX\_S8\_L002\_val\_2 |  |  |  |  |  | 74.4% | 49% | 134 bp | 9% | 53.6 |
| LNET10Tp4\_count.txt | 82.2% | 81.0 |  |  |  |  |  |  |  |  |
| LNET13TAHMVVKDRXX\_S19\_L001\_val\_1 |  |  |  |  |  | 79.9% | 50% | 135 bp | 18% | 52.9 |
| LNET13TAHMVVKDRXX\_S19\_L001\_val\_2 |  |  |  |  |  | 70.7% | 49% | 135 bp | 9% | 52.9 |
| LNET13TAHMVVKDRXX\_S19\_L002\_val\_1 |  |  |  |  |  | 79.4% | 50% | 135 bp | 18% | 53.2 |
| LNET13TAHMVVKDRXX\_S19\_L002\_val\_2 |  |  |  |  |  | 69.9% | 49% | 135 bp | 9% | 53.2 |
| LNET13T\_count.txt | 80.6% | 78.1 |  |  |  |  |  |  |  |  |
| LNET13Tp1AHMVVKDRXX\_S18\_L001\_val\_1 |  |  |  |  |  | 80.3% | 50% | 136 bp | 18% | 42.1 |
| LNET13Tp1AHMVVKDRXX\_S18\_L001\_val\_2 |  |  |  |  |  | 70.7% | 49% | 136 bp | 9% | 42.1 |
| LNET13Tp1AHMVVKDRXX\_S18\_L002\_val\_1 |  |  |  |  |  | 79.7% | 50% | 136 bp | 18% | 42.6 |
| LNET13Tp1AHMVVKDRXX\_S18\_L002\_val\_2 |  |  |  |  |  | 70.1% | 49% | 136 bp | 9% | 42.6 |
| LNET13Tp1\_count.txt | 80.5% | 62.1 |  |  |  |  |  |  |  |  |
| LNET14TAHMVVKDRXX\_S21\_L001\_val\_1 |  |  |  |  |  | 86.7% | 51% | 136 bp | 18% | 35.6 |
| LNET14TAHMVVKDRXX\_S21\_L001\_val\_2 |  |  |  |  |  | 78.5% | 50% | 136 bp | 9% | 35.6 |
| LNET14TAHMVVKDRXX\_S21\_L002\_val\_1 |  |  |  |  |  | 86.3% | 51% | 136 bp | 18% | 35.9 |
| LNET14TAHMVVKDRXX\_S21\_L002\_val\_2 |  |  |  |  |  | 77.9% | 50% | 136 bp | 9% | 35.9 |
| LNET14T\_count.txt | 73.5% | 47.9 |  |  |  |  |  |  |  |  |
| LNET14Tp1AHMVVKDRXX\_S20\_L001\_val\_1 |  |  |  |  |  | 80.5% | 51% | 137 bp | 18% | 42.9 |
| LNET14Tp1AHMVVKDRXX\_S20\_L001\_val\_2 |  |  |  |  |  | 69.5% | 50% | 137 bp | 9% | 42.9 |
| LNET14Tp1AHMVVKDRXX\_S20\_L002\_val\_1 |  |  |  |  |  | 80.1% | 51% | 137 bp | 18% | 43.3 |
| LNET14Tp1AHMVVKDRXX\_S20\_L002\_val\_2 |  |  |  |  |  | 69.2% | 50% | 137 bp | 9% | 43.3 |
| LNET14Tp1\_count.txt | 81.7% | 62.3 |  |  |  |  |  |  |  |  |
| LNET5TAHMVVKDRXX\_S3\_L001\_val\_1 |  |  |  |  |  | 79.1% | 51% | 136 bp | 18% | 50.2 |
| LNET5TAHMVVKDRXX\_S3\_L001\_val\_2 |  |  |  |  |  | 70.7% | 50% | 136 bp | 9% | 50.2 |
| LNET5TAHMVVKDRXX\_S3\_L002\_val\_1 |  |  |  |  |  | 78.9% | 51% | 137 bp | 18% | 50.9 |
| LNET5TAHMVVKDRXX\_S3\_L002\_val\_2 |  |  |  |  |  | 70.5% | 50% | 137 bp | 9% | 50.9 |
| LNET5T\_count.txt | 79.1% | 75.4 |  |  |  |  |  |  |  |  |
| LNET5Tp4AHMVVKDRXX\_S2\_L001\_val\_1 |  |  |  |  |  | 82.6% | 51% | 134 bp | 18% | 55.6 |
| LNET5Tp4AHMVVKDRXX\_S2\_L001\_val\_2 |  |  |  |  |  | 75.4% | 50% | 134 bp | 9% | 55.6 |
| LNET5Tp4AHMVVKDRXX\_S2\_L002\_val\_1 |  |  |  |  |  | 82.3% | 51% | 134 bp | 18% | 56.6 |
| LNET5Tp4AHMVVKDRXX\_S2\_L002\_val\_2 |  |  |  |  |  | 74.9% | 50% | 134 bp | 9% | 56.6 |
| LNET5Tp4\_count.txt | 83.2% | 88.8 |  |  |  |  |  |  |  |  |
| LNET6TAHMVVKDRXX\_S5\_L001\_val\_1 |  |  |  |  |  | 78.1% | 51% | 136 bp | 18% | 52.8 |
| LNET6TAHMVVKDRXX\_S5\_L001\_val\_2 |  |  |  |  |  | 69.4% | 50% | 136 bp | 9% | 52.8 |
| LNET6TAHMVVKDRXX\_S5\_L002\_val\_1 |  |  |  |  |  | 78.0% | 51% | 137 bp | 18% | 53.6 |
| LNET6TAHMVVKDRXX\_S5\_L002\_val\_2 |  |  |  |  |  | 68.8% | 50% | 136 bp | 9% | 53.6 |
| LNET6T\_count.txt | 81.2% | 80.8 |  |  |  |  |  |  |  |  |
| LNET6Tp1AHMVVKDRXX\_S4\_L001\_val\_1 |  |  |  |  |  | 78.4% | 51% | 133 bp | 18% | 54.2 |
| LNET6Tp1AHMVVKDRXX\_S4\_L001\_val\_2 |  |  |  |  |  | 69.0% | 50% | 133 bp | 9% | 54.2 |
| LNET6Tp1AHMVVKDRXX\_S4\_L002\_val\_1 |  |  |  |  |  | 77.9% | 51% | 133 bp | 18% | 54.6 |
| LNET6Tp1AHMVVKDRXX\_S4\_L002\_val\_2 |  |  |  |  |  | 68.4% | 50% | 134 bp | 9% | 54.6 |
| LNET6Tp1\_count.txt | 81.5% | 81.3 |  |  |  |  |  |  |  |  |
| PANEC1THVYYHBGXB\_S1\_L001\_val\_1 |  |  |  |  |  | 50.2% | 49% | 138 bp | 18% | 13.7 |
| PANEC1THVYYHBGXB\_S1\_L001\_val\_2 |  |  |  |  |  | 39.5% | 49% | 137 bp | 9% | 13.7 |
| PANEC1THVYYHBGXB\_S1\_L002\_val\_1 |  |  |  |  |  | 47.3% | 49% | 138 bp | 9% | 13.7 |
| PANEC1THVYYHBGXB\_S1\_L002\_val\_2 |  |  |  |  |  | 36.5% | 49% | 137 bp | 9% | 13.7 |
| PANEC1THVYYHBGXB\_S1\_L003\_val\_1 |  |  |  |  |  | 47.2% | 49% | 138 bp | 9% | 12.9 |
| PANEC1THVYYHBGXB\_S1\_L003\_val\_2 |  |  |  |  |  | 35.4% | 49% | 137 bp | 9% | 12.9 |
| PANEC1THVYYHBGXB\_S1\_L004\_val\_1 |  |  |  |  |  | 47.3% | 49% | 138 bp | 9% | 12.9 |
| PANEC1THVYYHBGXB\_S1\_L004\_val\_2 |  |  |  |  |  | 36.0% | 49% | 137 bp | 9% | 12.9 |
| PANEC1T\_count.txt | 78.1% | 35.8 |  |  |  |  |  |  |  |  |
| PANEC1Tp14HVYYHBGXB\_S6\_L001\_val\_1 |  |  |  |  |  | 41.0% | 48% | 137 bp | 9% | 7.9 |
| PANEC1Tp14HVYYHBGXB\_S6\_L001\_val\_2 |  |  |  |  |  | 30.7% | 49% | 136 bp | 9% | 7.9 |
| PANEC1Tp14HVYYHBGXB\_S6\_L002\_val\_1 |  |  |  |  |  | 39.0% | 48% | 137 bp | 9% | 7.8 |
| PANEC1Tp14HVYYHBGXB\_S6\_L002\_val\_2 |  |  |  |  |  | 27.7% | 49% | 136 bp | 9% | 7.8 |
| PANEC1Tp14HVYYHBGXB\_S6\_L003\_val\_1 |  |  |  |  |  | 39.0% | 48% | 137 bp | 9% | 7.4 |
| PANEC1Tp14HVYYHBGXB\_S6\_L003\_val\_2 |  |  |  |  |  | 27.6% | 49% | 136 bp | 9% | 7.4 |
| PANEC1Tp14HVYYHBGXB\_S6\_L004\_val\_1 |  |  |  |  |  | 39.0% | 48% | 137 bp | 9% | 7.3 |
| PANEC1Tp14HVYYHBGXB\_S6\_L004\_val\_2 |  |  |  |  |  | 28.1% | 49% | 136 bp | 9% | 7.3 |
| PANEC1Tp14\_count.txt | 81.1% | 21.5 |  |  |  |  |  |  |  |  |
| PANEC1Tp4HVYYHBGXB\_S7\_L001\_val\_1 |  |  |  |  |  | 53.5% | 48% | 136 bp | 18% | 15.9 |
| PANEC1Tp4HVYYHBGXB\_S7\_L001\_val\_2 |  |  |  |  |  | 42.6% | 48% | 136 bp | 9% | 15.9 |
| PANEC1Tp4HVYYHBGXB\_S7\_L002\_val\_1 |  |  |  |  |  | 50.9% | 48% | 137 bp | 18% | 15.7 |
| PANEC1Tp4HVYYHBGXB\_S7\_L002\_val\_2 |  |  |  |  |  | 38.7% | 48% | 136 bp | 9% | 15.7 |
| PANEC1Tp4HVYYHBGXB\_S7\_L003\_val\_1 |  |  |  |  |  | 50.5% | 48% | 136 bp | 18% | 14.8 |
| PANEC1Tp4HVYYHBGXB\_S7\_L003\_val\_2 |  |  |  |  |  | 37.8% | 48% | 136 bp | 9% | 14.8 |
| PANEC1Tp4HVYYHBGXB\_S7\_L004\_val\_1 |  |  |  |  |  | 50.3% | 48% | 137 bp | 18% | 14.7 |
| PANEC1Tp4HVYYHBGXB\_S7\_L004\_val\_2 |  |  |  |  |  | 38.3% | 48% | 136 bp | 9% | 14.7 |
| PANEC1Tp4\_count.txt | 80.3% | 43.6 |  |  |  |  |  |  |  |  |
| SINET12MAHMVVKDRXX\_S16\_L001\_val\_1 |  |  |  |  |  | 85.7% | 50% | 132 bp | 18% | 38.0 |
| SINET12MAHMVVKDRXX\_S16\_L001\_val\_2 |  |  |  |  |  | 75.6% | 49% | 132 bp | 18% | 38.0 |
| SINET12MAHMVVKDRXX\_S16\_L002\_val\_1 |  |  |  |  |  | 85.3% | 50% | 132 bp | 18% | 38.2 |
| SINET12MAHMVVKDRXX\_S16\_L002\_val\_2 |  |  |  |  |  | 75.0% | 49% | 133 bp | 18% | 38.2 |
| SINET12M\_count.txt | 79.8% | 53.9 |  |  |  |  |  |  |  |  |
| SINET12Mp1.1AHMVVKDRXX\_S17\_L001\_val\_1 |  |  |  |  |  | 76.6% | 51% | 136 bp | 18% | 47.3 |
| SINET12Mp1.1AHMVVKDRXX\_S17\_L001\_val\_2 |  |  |  |  |  | 67.4% | 50% | 136 bp | 9% | 47.3 |
| SINET12Mp1.1AHMVVKDRXX\_S17\_L002\_val\_1 |  |  |  |  |  | 76.1% | 51% | 136 bp | 18% | 47.9 |
| SINET12Mp1.1AHMVVKDRXX\_S17\_L002\_val\_2 |  |  |  |  |  | 66.5% | 50% | 136 bp | 9% | 47.9 |
| SINET12Mp1.1\_count.txt | 82.7% | 74.4 |  |  |  |  |  |  |  |  |
| SINET12Mp1.3AHMVVKDRXX\_S15\_L001\_val\_1 |  |  |  |  |  | 77.4% | 51% | 135 bp | 18% | 36.9 |
| SINET12Mp1.3AHMVVKDRXX\_S15\_L001\_val\_2 |  |  |  |  |  | 65.8% | 50% | 135 bp | 9% | 36.9 |
| SINET12Mp1.3AHMVVKDRXX\_S15\_L002\_val\_1 |  |  |  |  |  | 77.0% | 51% | 136 bp | 18% | 37.2 |
| SINET12Mp1.3AHMVVKDRXX\_S15\_L002\_val\_2 |  |  |  |  |  | 65.3% | 50% | 135 bp | 9% | 37.2 |
| SINET12Mp1.3\_count.txt | 84.5% | 56.3 |  |  |  |  |  |  |  |  |
| SINET7MAHMVVKDRXX\_S11\_L001\_val\_1 |  |  |  |  |  | 79.0% | 51% | 136 bp | 18% | 33.0 |
| SINET7MAHMVVKDRXX\_S11\_L001\_val\_2 |  |  |  |  |  | 66.0% | 50% | 135 bp | 18% | 33.0 |
| SINET7MAHMVVKDRXX\_S11\_L002\_val\_1 |  |  |  |  |  | 78.8% | 52% | 136 bp | 18% | 33.4 |
| SINET7MAHMVVKDRXX\_S11\_L002\_val\_2 |  |  |  |  |  | 65.7% | 50% | 136 bp | 18% | 33.4 |
| SINET7M\_count.txt | 82.5% | 47.0 |  |  |  |  |  |  |  |  |
| SINET7Mp2AHMVVKDRXX\_S10\_L001\_val\_1 |  |  |  |  |  | 77.0% | 51% | 134 bp | 18% | 44.0 |
| SINET7Mp2AHMVVKDRXX\_S10\_L001\_val\_2 |  |  |  |  |  | 67.4% | 50% | 134 bp | 9% | 44.0 |
| SINET7Mp2AHMVVKDRXX\_S10\_L002\_val\_1 |  |  |  |  |  | 76.8% | 51% | 134 bp | 18% | 44.5 |
| SINET7Mp2AHMVVKDRXX\_S10\_L002\_val\_2 |  |  |  |  |  | 67.1% | 50% | 134 bp | 9% | 44.5 |
| SINET7Mp2\_count.txt | 82.7% | 67.8 |  |  |  |  |  |  |  |  |
| SINET8MAHMVVKDRXX\_S13\_L001\_val\_1 |  |  |  |  |  | 78.1% | 52% | 136 bp | 18% | 44.1 |
| SINET8MAHMVVKDRXX\_S13\_L001\_val\_2 |  |  |  |  |  | 69.2% | 51% | 136 bp | 9% | 44.1 |
| SINET8MAHMVVKDRXX\_S13\_L002\_val\_1 |  |  |  |  |  | 77.8% | 52% | 136 bp | 18% | 44.5 |
| SINET8MAHMVVKDRXX\_S13\_L002\_val\_2 |  |  |  |  |  | 68.8% | 51% | 136 bp | 9% | 44.5 |
| SINET8M\_count.txt | 82.0% | 68.2 |  |  |  |  |  |  |  |  |
| SINET8Mp2AHMVVKDRXX\_S12\_L001\_val\_1 |  |  |  |  |  | 77.8% | 51% | 133 bp | 18% | 46.2 |
| SINET8Mp2AHMVVKDRXX\_S12\_L001\_val\_2 |  |  |  |  |  | 69.0% | 51% | 133 bp | 9% | 46.2 |
| SINET8Mp2AHMVVKDRXX\_S12\_L002\_val\_1 |  |  |  |  |  | 77.0% | 51% | 133 bp | 18% | 46.5 |
| SINET8Mp2AHMVVKDRXX\_S12\_L002\_val\_2 |  |  |  |  |  | 68.2% | 51% | 133 bp | 9% | 46.5 |
| SINET8Mp2\_count.txt | 84.3% | 73.4 |  |  |  |  |  |  |  |  |
| STAR.LCNEC11M.Log.final.out |  |  | 87.5% | 88.2 |  |  |  |  |  |  |
| STAR.LCNEC11Mp3.Log.final.out |  |  | 85.6% | 82.6 |  |  |  |  |  |  |
| STAR.LCNEC3T.Log.final.out |  |  | 75.6% | 42.4 |  |  |  |  |  |  |
| STAR.LCNEC3Tp17.2.Log.final.out |  |  | 85.2% | 88.2 |  |  |  |  |  |  |
| STAR.LCNEC3Tp17.Log.final.out |  |  | 81.2% | 35.3 |  |  |  |  |  |  |
| STAR.LCNEC4T.Log.final.out |  |  | 78.7% | 44.4 |  |  |  |  |  |  |
| STAR.LCNEC4Tp24.Log.final.out |  |  | 81.8% | 42.0 |  |  |  |  |  |  |
| STAR.LCNEC4Tp7.Log.final.out |  |  | 84.3% | 76.7 |  |  |  |  |  |  |
| STAR.LNET10T.Log.final.out |  |  | 76.6% | 64.4 |  |  |  |  |  |  |
| STAR.LNET10Tp4.Log.final.out |  |  | 87.2% | 93.0 |  |  |  |  |  |  |
| STAR.LNET13T.Log.final.out |  |  | 84.9% | 90.1 |  |  |  |  |  |  |
| STAR.LNET13Tp1.Log.final.out |  |  | 84.1% | 71.3 |  |  |  |  |  |  |
| STAR.LNET14T.Log.final.out |  |  | 82.1% | 58.8 |  |  |  |  |  |  |
| STAR.LNET14Tp1.Log.final.out |  |  | 82.3% | 71.0 |  |  |  |  |  |  |
| STAR.LNET5T.Log.final.out |  |  | 87.2% | 88.1 |  |  |  |  |  |  |
| STAR.LNET5Tp4.Log.final.out |  |  | 88.6% | 99.4 |  |  |  |  |  |  |
| STAR.LNET6T.Log.final.out |  |  | 87.6% | 93.2 |  |  |  |  |  |  |
| STAR.LNET6Tp1.Log.final.out |  |  | 87.2% | 94.9 |  |  |  |  |  |  |
| STAR.PANEC1T.Log.final.out |  |  | 79.1% | 42.0 |  |  |  |  |  |  |
| STAR.PANEC1Tp14.Log.final.out |  |  | 81.7% | 24.9 |  |  |  |  |  |  |
| STAR.PANEC1Tp4.Log.final.out |  |  | 80.9% | 49.4 |  |  |  |  |  |  |
| STAR.SINET12M.Log.final.out |  |  | 82.3% | 62.7 |  |  |  |  |  |  |
| STAR.SINET12Mp1.1.Log.final.out |  |  | 90.0% | 85.6 |  |  |  |  |  |  |
| STAR.SINET12Mp1.3.Log.final.out |  |  | 86.1% | 63.7 |  |  |  |  |  |  |
| STAR.SINET7M.Log.final.out |  |  | 79.5% | 52.8 |  |  |  |  |  |  |
| STAR.SINET7Mp2.Log.final.out |  |  | 88.2% | 78.1 |  |  |  |  |  |  |
| STAR.SINET8M.Log.final.out |  |  | 87.9% | 77.9 |  |  |  |  |  |  |
| STAR.SINET8Mp2.Log.final.out |  |  | 89.4% | 82.9 |  |  |  |  |  |  |
| TD10B\_AHMVVKDRXX\_S8\_L001\_R1\_001 |  |  |  |  | 11.8% |  |  |  |  |  |
| TD10B\_AHMVVKDRXX\_S8\_L001\_R2\_001 |  |  |  |  | 11.6% |  |  |  |  |  |
| TD10B\_AHMVVKDRXX\_S8\_L002\_R1\_001 |  |  |  |  | 11.7% |  |  |  |  |  |
| TD10B\_AHMVVKDRXX\_S8\_L002\_R2\_001 |  |  |  |  | 11.5% |  |  |  |  |  |
| TD10C\_AHMVVKDRXX\_S9\_L001\_R1\_001 |  |  |  |  | 12.5% |  |  |  |  |  |
| TD10C\_AHMVVKDRXX\_S9\_L001\_R2\_001 |  |  |  |  | 13.4% |  |  |  |  |  |
| TD10C\_AHMVVKDRXX\_S9\_L002\_R1\_001 |  |  |  |  | 12.3% |  |  |  |  |  |
| TD10C\_AHMVVKDRXX\_S9\_L002\_R2\_001 |  |  |  |  | 13.3% |  |  |  |  |  |
| TD11B\_AHMVVKDRXX\_S6\_L001\_R1\_001 |  |  |  |  | 12.2% |  |  |  |  |  |
| TD11B\_AHMVVKDRXX\_S6\_L001\_R2\_001 |  |  |  |  | 12.2% |  |  |  |  |  |
| TD11B\_AHMVVKDRXX\_S6\_L002\_R1\_001 |  |  |  |  | 12.0% |  |  |  |  |  |
| TD11B\_AHMVVKDRXX\_S6\_L002\_R2\_001 |  |  |  |  | 12.0% |  |  |  |  |  |
| TD11C\_AHMVVKDRXX\_S7\_L001\_R1\_001 |  |  |  |  | 10.9% |  |  |  |  |  |
| TD11C\_AHMVVKDRXX\_S7\_L001\_R2\_001 |  |  |  |  | 11.3% |  |  |  |  |  |
| TD11C\_AHMVVKDRXX\_S7\_L002\_R1\_001 |  |  |  |  | 10.8% |  |  |  |  |  |
| TD11C\_AHMVVKDRXX\_S7\_L002\_R2\_001 |  |  |  |  | 11.2% |  |  |  |  |  |
| TD12B\_AHMVVKDRXX\_S15\_L001\_R1\_001 |  |  |  |  | 10.5% |  |  |  |  |  |
| TD12B\_AHMVVKDRXX\_S15\_L001\_R2\_001 |  |  |  |  | 11.3% |  |  |  |  |  |
| TD12B\_AHMVVKDRXX\_S15\_L002\_R1\_001 |  |  |  |  | 10.4% |  |  |  |  |  |
| TD12B\_AHMVVKDRXX\_S15\_L002\_R2\_001 |  |  |  |  | 11.2% |  |  |  |  |  |
| TD12C\_AHMVVKDRXX\_S16\_L001\_R1\_001 |  |  |  |  | 13.0% |  |  |  |  |  |
| TD12C\_AHMVVKDRXX\_S16\_L001\_R2\_001 |  |  |  |  | 13.1% |  |  |  |  |  |
| TD12C\_AHMVVKDRXX\_S16\_L002\_R1\_001 |  |  |  |  | 12.8% |  |  |  |  |  |
| TD12C\_AHMVVKDRXX\_S16\_L002\_R2\_001 |  |  |  |  | 12.9% |  |  |  |  |  |
| TD12D\_AHMVVKDRXX\_S17\_L001\_R1\_001 |  |  |  |  | 10.4% |  |  |  |  |  |
| TD12D\_AHMVVKDRXX\_S17\_L001\_R2\_001 |  |  |  |  | 10.3% |  |  |  |  |  |
| TD12D\_AHMVVKDRXX\_S17\_L002\_R1\_001 |  |  |  |  | 10.3% |  |  |  |  |  |
| TD12D\_AHMVVKDRXX\_S17\_L002\_R2\_001 |  |  |  |  | 10.2% |  |  |  |  |  |
| TD13B\_AHMVVKDRXX\_S18\_L001\_R1\_001 |  |  |  |  | 10.4% |  |  |  |  |  |
| TD13B\_AHMVVKDRXX\_S18\_L001\_R2\_001 |  |  |  |  | 10.7% |  |  |  |  |  |
| TD13B\_AHMVVKDRXX\_S18\_L002\_R1\_001 |  |  |  |  | 10.3% |  |  |  |  |  |
| TD13B\_AHMVVKDRXX\_S18\_L002\_R2\_001 |  |  |  |  | 10.6% |  |  |  |  |  |
| TD13C\_AHMVVKDRXX\_S19\_L001\_R1\_001 |  |  |  |  | 10.9% |  |  |  |  |  |
| TD13C\_AHMVVKDRXX\_S19\_L001\_R2\_001 |  |  |  |  | 11.0% |  |  |  |  |  |
| TD13C\_AHMVVKDRXX\_S19\_L002\_R1\_001 |  |  |  |  | 10.7% |  |  |  |  |  |
| TD13C\_AHMVVKDRXX\_S19\_L002\_R2\_001 |  |  |  |  | 10.9% |  |  |  |  |  |
| TD14B\_AHMVVKDRXX\_S20\_L001\_R1\_001 |  |  |  |  | 9.9% |  |  |  |  |  |
| TD14B\_AHMVVKDRXX\_S20\_L001\_R2\_001 |  |  |  |  | 10.3% |  |  |  |  |  |
| TD14B\_AHMVVKDRXX\_S20\_L002\_R1\_001 |  |  |  |  | 9.8% |  |  |  |  |  |
| TD14B\_AHMVVKDRXX\_S20\_L002\_R2\_001 |  |  |  |  | 10.1% |  |  |  |  |  |
| TD14C\_AHMVVKDRXX\_S21\_L001\_R1\_001 |  |  |  |  | 12.0% |  |  |  |  |  |
| TD14C\_AHMVVKDRXX\_S21\_L001\_R2\_001 |  |  |  |  | 10.2% |  |  |  |  |  |
| TD14C\_AHMVVKDRXX\_S21\_L002\_R1\_001 |  |  |  |  | 11.9% |  |  |  |  |  |
| TD14C\_AHMVVKDRXX\_S21\_L002\_R2\_001 |  |  |  |  | 10.0% |  |  |  |  |  |
| TD1A\_HVYYHBGXB\_S6\_L001\_R1\_001 |  |  |  |  | 0.9% |  |  |  |  |  |
| TD1A\_HVYYHBGXB\_S6\_L001\_R2\_001 |  |  |  |  | 12.2% |  |  |  |  |  |
| TD1A\_HVYYHBGXB\_S6\_L002\_R1\_001 |  |  |  |  | 0.9% |  |  |  |  |  |
| TD1A\_HVYYHBGXB\_S6\_L002\_R2\_001 |  |  |  |  | 12.1% |  |  |  |  |  |
| TD1A\_HVYYHBGXB\_S6\_L003\_R1\_001 |  |  |  |  | 1.1% |  |  |  |  |  |
| TD1A\_HVYYHBGXB\_S6\_L003\_R2\_001 |  |  |  |  | 12.1% |  |  |  |  |  |
| TD1A\_HVYYHBGXB\_S6\_L004\_R1\_001 |  |  |  |  | 1.1% |  |  |  |  |  |
| TD1A\_HVYYHBGXB\_S6\_L004\_R2\_001 |  |  |  |  | 12.3% |  |  |  |  |  |
| TD1B\_HVYYHBGXB\_S7\_L001\_R1\_001 |  |  |  |  | 0.9% |  |  |  |  |  |
| TD1B\_HVYYHBGXB\_S7\_L001\_R2\_001 |  |  |  |  | 11.6% |  |  |  |  |  |
| TD1B\_HVYYHBGXB\_S7\_L002\_R1\_001 |  |  |  |  | 0.9% |  |  |  |  |  |
| TD1B\_HVYYHBGXB\_S7\_L002\_R2\_001 |  |  |  |  | 11.6% |  |  |  |  |  |
| TD1B\_HVYYHBGXB\_S7\_L003\_R1\_001 |  |  |  |  | 1.0% |  |  |  |  |  |
| TD1B\_HVYYHBGXB\_S7\_L003\_R2\_001 |  |  |  |  | 11.5% |  |  |  |  |  |
| TD1B\_HVYYHBGXB\_S7\_L004\_R1\_001 |  |  |  |  | 1.0% |  |  |  |  |  |
| TD1B\_HVYYHBGXB\_S7\_L004\_R2\_001 |  |  |  |  | 11.6% |  |  |  |  |  |
| TD1C\_HVYYHBGXB\_S1\_L001\_R1\_001 |  |  |  |  | 1.0% |  |  |  |  |  |
| TD1C\_HVYYHBGXB\_S1\_L001\_R2\_001 |  |  |  |  | 11.9% |  |  |  |  |  |
| TD1C\_HVYYHBGXB\_S1\_L002\_R1\_001 |  |  |  |  | 0.9% |  |  |  |  |  |
| TD1C\_HVYYHBGXB\_S1\_L002\_R2\_001 |  |  |  |  | 11.8% |  |  |  |  |  |
| TD1C\_HVYYHBGXB\_S1\_L003\_R1\_001 |  |  |  |  | 1.1% |  |  |  |  |  |
| TD1C\_HVYYHBGXB\_S1\_L003\_R2\_001 |  |  |  |  | 11.8% |  |  |  |  |  |
| TD1C\_HVYYHBGXB\_S1\_L004\_R1\_001 |  |  |  |  | 1.1% |  |  |  |  |  |
| TD1C\_HVYYHBGXB\_S1\_L004\_R2\_001 |  |  |  |  | 11.9% |  |  |  |  |  |
| TD3A\_HVYYHBGXB\_S2\_L001\_R1\_001 |  |  |  |  | 0.9% |  |  |  |  |  |
| TD3A\_HVYYHBGXB\_S2\_L001\_R2\_001 |  |  |  |  | 10.0% |  |  |  |  |  |
| TD3A\_HVYYHBGXB\_S2\_L002\_R1\_001 |  |  |  |  | 0.8% |  |  |  |  |  |
| TD3A\_HVYYHBGXB\_S2\_L002\_R2\_001 |  |  |  |  | 9.9% |  |  |  |  |  |
| TD3A\_HVYYHBGXB\_S2\_L003\_R1\_001 |  |  |  |  | 1.0% |  |  |  |  |  |
| TD3A\_HVYYHBGXB\_S2\_L003\_R2\_001 |  |  |  |  | 9.9% |  |  |  |  |  |
| TD3A\_HVYYHBGXB\_S2\_L004\_R1\_001 |  |  |  |  | 1.0% |  |  |  |  |  |
| TD3A\_HVYYHBGXB\_S2\_L004\_R2\_001 |  |  |  |  | 10.0% |  |  |  |  |  |
| TD3B\_HVYYHBGXB\_S3\_L001\_R1\_001 |  |  |  |  | 0.9% |  |  |  |  |  |
| TD3B\_HVYYHBGXB\_S3\_L001\_R2\_001 |  |  |  |  | 11.3% |  |  |  |  |  |
| TD3B\_HVYYHBGXB\_S3\_L002\_R1\_001 |  |  |  |  | 0.9% |  |  |  |  |  |
| TD3B\_HVYYHBGXB\_S3\_L002\_R2\_001 |  |  |  |  | 11.2% |  |  |  |  |  |
| TD3B\_HVYYHBGXB\_S3\_L003\_R1\_001 |  |  |  |  | 1.1% |  |  |  |  |  |
| TD3B\_HVYYHBGXB\_S3\_L003\_R2\_001 |  |  |  |  | 11.2% |  |  |  |  |  |
| TD3B\_HVYYHBGXB\_S3\_L004\_R1\_001 |  |  |  |  | 1.1% |  |  |  |  |  |
| TD3B\_HVYYHBGXB\_S3\_L004\_R2\_001 |  |  |  |  | 11.3% |  |  |  |  |  |
| TD3E\_AHMVVKDRXX\_S1\_L001\_R1\_001 |  |  |  |  | 13.7% |  |  |  |  |  |
| TD3E\_AHMVVKDRXX\_S1\_L001\_R2\_001 |  |  |  |  | 14.1% |  |  |  |  |  |
| TD3E\_AHMVVKDRXX\_S1\_L002\_R1\_001 |  |  |  |  | 13.6% |  |  |  |  |  |
| TD3E\_AHMVVKDRXX\_S1\_L002\_R2\_001 |  |  |  |  | 14.0% |  |  |  |  |  |
| TD4A\_HVYYHBGXB\_S5\_L001\_R1\_001 |  |  |  |  | 0.9% |  |  |  |  |  |
| TD4A\_HVYYHBGXB\_S5\_L001\_R2\_001 |  |  |  |  | 11.3% |  |  |  |  |  |
| TD4A\_HVYYHBGXB\_S5\_L002\_R1\_001 |  |  |  |  | 0.9% |  |  |  |  |  |
| TD4A\_HVYYHBGXB\_S5\_L002\_R2\_001 |  |  |  |  | 11.2% |  |  |  |  |  |
| TD4A\_HVYYHBGXB\_S5\_L003\_R1\_001 |  |  |  |  | 1.0% |  |  |  |  |  |
| TD4A\_HVYYHBGXB\_S5\_L003\_R2\_001 |  |  |  |  | 11.2% |  |  |  |  |  |
| TD4A\_HVYYHBGXB\_S5\_L004\_R1\_001 |  |  |  |  | 1.0% |  |  |  |  |  |
| TD4A\_HVYYHBGXB\_S5\_L004\_R2\_001 |  |  |  |  | 11.3% |  |  |  |  |  |
| TD4D\_AHMVVKDRXX\_S14\_L001\_R1\_001 |  |  |  |  | 12.5% |  |  |  |  |  |
| TD4D\_AHMVVKDRXX\_S14\_L001\_R2\_001 |  |  |  |  | 12.5% |  |  |  |  |  |
| TD4D\_AHMVVKDRXX\_S14\_L002\_R1\_001 |  |  |  |  | 12.3% |  |  |  |  |  |
| TD4D\_AHMVVKDRXX\_S14\_L002\_R2\_001 |  |  |  |  | 12.4% |  |  |  |  |  |
| TD4\_HVYYHBGXB\_S4\_L001\_R1\_001 |  |  |  |  | 0.8% |  |  |  |  |  |
| TD4\_HVYYHBGXB\_S4\_L001\_R2\_001 |  |  |  |  | 10.2% |  |  |  |  |  |
| TD4\_HVYYHBGXB\_S4\_L002\_R1\_001 |  |  |  |  | 0.8% |  |  |  |  |  |
| TD4\_HVYYHBGXB\_S4\_L002\_R2\_001 |  |  |  |  | 10.1% |  |  |  |  |  |
| TD4\_HVYYHBGXB\_S4\_L003\_R1\_001 |  |  |  |  | 0.9% |  |  |  |  |  |
| TD4\_HVYYHBGXB\_S4\_L003\_R2\_001 |  |  |  |  | 10.1% |  |  |  |  |  |
| TD4\_HVYYHBGXB\_S4\_L004\_R1\_001 |  |  |  |  | 0.9% |  |  |  |  |  |
| TD4\_HVYYHBGXB\_S4\_L004\_R2\_001 |  |  |  |  | 10.2% |  |  |  |  |  |
| TD5B\_AHMVVKDRXX\_S2\_L001\_R1\_001 |  |  |  |  | 11.3% |  |  |  |  |  |
| TD5B\_AHMVVKDRXX\_S2\_L001\_R2\_001 |  |  |  |  | 11.4% |  |  |  |  |  |
| TD5B\_AHMVVKDRXX\_S2\_L002\_R1\_001 |  |  |  |  | 11.2% |  |  |  |  |  |
| TD5B\_AHMVVKDRXX\_S2\_L002\_R2\_001 |  |  |  |  | 11.2% |  |  |  |  |  |
| TD5C\_AHMVVKDRXX\_S3\_L001\_R1\_001 |  |  |  |  | 10.1% |  |  |  |  |  |
| TD5C\_AHMVVKDRXX\_S3\_L001\_R2\_001 |  |  |  |  | 10.0% |  |  |  |  |  |
| TD5C\_AHMVVKDRXX\_S3\_L002\_R1\_001 |  |  |  |  | 10.0% |  |  |  |  |  |
| TD5C\_AHMVVKDRXX\_S3\_L002\_R2\_001 |  |  |  |  | 9.9% |  |  |  |  |  |
| TD6B\_AHMVVKDRXX\_S4\_L001\_R1\_001 |  |  |  |  | 11.9% |  |  |  |  |  |
| TD6B\_AHMVVKDRXX\_S4\_L001\_R2\_001 |  |  |  |  | 12.2% |  |  |  |  |  |
| TD6B\_AHMVVKDRXX\_S4\_L002\_R1\_001 |  |  |  |  | 11.8% |  |  |  |  |  |
| TD6B\_AHMVVKDRXX\_S4\_L002\_R2\_001 |  |  |  |  | 12.0% |  |  |  |  |  |
| TD6C\_AHMVVKDRXX\_S5\_L001\_R1\_001 |  |  |  |  | 10.3% |  |  |  |  |  |
| TD6C\_AHMVVKDRXX\_S5\_L001\_R2\_001 |  |  |  |  | 10.1% |  |  |  |  |  |
| TD6C\_AHMVVKDRXX\_S5\_L002\_R1\_001 |  |  |  |  | 10.1% |  |  |  |  |  |
| TD6C\_AHMVVKDRXX\_S5\_L002\_R2\_001 |  |  |  |  | 10.0% |  |  |  |  |  |
| TD7B\_AHMVVKDRXX\_S10\_L001\_R1\_001 |  |  |  |  | 11.7% |  |  |  |  |  |
| TD7B\_AHMVVKDRXX\_S10\_L001\_R2\_001 |  |  |  |  | 11.7% |  |  |  |  |  |
| TD7B\_AHMVVKDRXX\_S10\_L002\_R1\_001 |  |  |  |  | 11.5% |  |  |  |  |  |
| TD7B\_AHMVVKDRXX\_S10\_L002\_R2\_001 |  |  |  |  | 11.5% |  |  |  |  |  |
| TD7C\_AHMVVKDRXX\_S11\_L001\_R1\_001 |  |  |  |  | 10.8% |  |  |  |  |  |
| TD7C\_AHMVVKDRXX\_S11\_L001\_R2\_001 |  |  |  |  | 11.4% |  |  |  |  |  |
| TD7C\_AHMVVKDRXX\_S11\_L002\_R1\_001 |  |  |  |  | 10.6% |  |  |  |  |  |
| TD7C\_AHMVVKDRXX\_S11\_L002\_R2\_001 |  |  |  |  | 11.2% |  |  |  |  |  |
| TD8B\_AHMVVKDRXX\_S12\_L001\_R1\_001 |  |  |  |  | 12.5% |  |  |  |  |  |
| TD8B\_AHMVVKDRXX\_S12\_L001\_R2\_001 |  |  |  |  | 12.4% |  |  |  |  |  |
| TD8B\_AHMVVKDRXX\_S12\_L002\_R1\_001 |  |  |  |  | 12.3% |  |  |  |  |  |
| TD8B\_AHMVVKDRXX\_S12\_L002\_R2\_001 |  |  |  |  | 12.3% |  |  |  |  |  |
| TD8C\_AHMVVKDRXX\_S13\_L001\_R1\_001 |  |  |  |  | 10.1% |  |  |  |  |  |
| TD8C\_AHMVVKDRXX\_S13\_L001\_R2\_001 |  |  |  |  | 10.3% |  |  |  |  |  |
| TD8C\_AHMVVKDRXX\_S13\_L002\_R1\_001 |  |  |  |  | 9.9% |  |  |  |  |  |
| TD8C\_AHMVVKDRXX\_S13\_L002\_R2\_001 |  |  |  |  | 10.2% |  |  |  |  |  |

×

###### General Statistics: Columns

Uncheck the tick box to hide columns. Click and drag the handle on the left to change order.

Show All
Show None

| Sort | Visible | Group | Column | Description | ID | Scale |
| --- | --- | --- | --- | --- | --- | --- |
| \|\| |  | HTSeq Count | % Assigned | % Assigned reads | `percent_assigned` | None |
| \|\| |  | HTSeq Count | M Assigned | Assigned Reads (millions) | `assigned` | read\_count |
| \|\| |  | STAR | % Aligned | % Uniquely mapped reads | `uniquely_mapped_percent` | None |
| \|\| |  | STAR | M Aligned | Uniquely mapped reads (millions) | `uniquely_mapped` | read\_count |
| \|\| |  | Cutadapt | % Trimmed | % Total Base Pairs trimmed | `percent_trimmed` | None |
| \|\| |  | FastQC | % Dups | % Duplicate Reads | `percent_duplicates` | None |
| \|\| |  | FastQC | % GC | Average % GC Content | `percent_gc` | None |
| \|\| |  | FastQC | Length | Average Sequence Length (bp) | `avg_sequence_length` | None |
| \|\| |  | FastQC | % Failed | Percentage of modules failed in FastQC report (includes those not plotted here) | `percent_fails` | None |
| \|\| |  | FastQC | M Seqs | Total Sequences (millions) | `total_sequences` | read\_count |

Close

#### RSeQC

RSeQC package provides a number of useful modules that can comprehensively evaluate high throughput RNA-seq data.

##### Read Distribution

Read Distribution calculates how mapped reads are distributed over genome features.

Number of Tags
Percentages

loading..

---

##### Junction Saturation

Junction Saturation
counts the number of known splicing junctions that are observed
in each dataset. If sequencing depth is sufficient, all (annotated) splice junctions should
be rediscovered, resulting in a curve that reaches a plateau. Missing low abundance splice
junctions can affect downstream analysis.

Click a line to see the data side by side (as in the original RSeQC plot).

All Junctions
Known Junctions
Novel Junctions

loading..

---

#### HTSeq Count

HTSeq Count is part of the HTSeq Python package - it takes a file with aligned sequencing reads, plus a list of genomic features and counts how many reads map to each feature.

Number of Reads
Percentages

loading..

---

#### STAR

STAR is an ultrafast universal RNA-seq aligner.

##### Alignment Scores

Number of Reads
Percentages

loading..

---

#### Cutadapt

Cutadapt is a tool to find and remove adapter sequences, primers, poly-Atails and other types of unwanted sequence from your high-throughput sequencing reads.

This plot shows the number of reads with certain lengths of adapter trimmed.
Obs/Exp shows the raw counts divided by the number expected due to sequencing errors. A defined peak
may be related to adapter length. See the
cutadapt documentation
for more information on how these numbers are generated.

Click a sample row to see a line plot for that dataset.

###### Rollover for sample name

 Export Plot

Position: -

%T: -

%C: -

%A: -

%G: -

---

##### Per Sequence GC Content Help

**The dashed black line shows theoretical GC content:** `Human Transcriptome (UCSC hg38)`

Sort by highlight

loading..

**MultiQC v1.8** 
- Written by Phil Ewels,
available on GitHub.

This report uses HighCharts,
jQuery,
jQuery UI,
Bootstrap,
FileSaver.js and
clipboard.js.

×

##### Plot Table Data

Select Column

Select Column

Please select two table columns.

Close

×

##### Regex Help

Toolbox search strings can behave as regular expressions (regexes). Click a button below to see an example of it in action. Try modifying them yourself in the text box.

`^` (start of string)
`$` (end of string)
`[]` (character choice)
`\d` (shorthand for `[0-9]`)
`\w` (shorthand for `[0-9a-zA-Z_]`)
`.` (any character)
`\.` (literal full stop)
`()` `|` (group / separator)
`*` (prev char 0 or more)
`+` (prev char 1 or more)
`?` (prev char 0 or 1)
`{}` (char num times)
`{,}` (count range)

```
samp_1
samp_1_edited
samp_2
samp_2_edited
samp_3
samp_3_edited
prepended_samp_1
tmp_samp_1_edited
tmpp_samp_1_edited
tmppp_samp_1_edited
#samp_1_edited.tmp
samp_11
samp_11111
```

See regex101.com for a more heavy duty testing suite.

Close
